# *In silico* screening and testing of FDA approved small molecules to block SARS-CoV-2 entry to the host cell by inhibiting Spike protein cleavage

**DOI:** 10.1101/2022.03.07.483324

**Authors:** E. Sila Ozdemir, Hillary H. Le, Adem Yildirim, Srivathsan V. Ranganathan

## Abstract

The COVID-19 pandemic began in 2019, but it is still active. The development of an effective vaccine reduced the number of deaths; however, a treatment is still needed. Here, we aimed to inhibit viral entry to the host cell by inhibiting Spike (S) protein cleavage by several proteases. We develop a computational pipeline to repurpose FDA-approved drugs to inhibit protease activity and thus prevent S protein cleavage. We tested some of our drug candidates and demonstrated a decrease in protease activity. We believe our pipeline will be beneficial in identifying a drug regimen for COVID-19 patients.

## 1 Introduction

The outbreak of the novel severe acute respiratory syndrome coronavirus 2 (SARS-CoV-2) in China is followed by a global level emergency. COVID-19, the disease caused by SARS-CoV-2, is a global pandemic, has resulted in over five million deaths, and the death toll continues to climb, which is devastating as the virus is expected to be in circulation for a long time (1). The novel coronavirus has been sequenced and it has been revealed that it shares substantial similarity with the SARS-CoV that caused a similar epidemic in 2003 (2), they are both zoonotic coronaviruses belonging to the betacoronavirus family (3).

The world is in search of a robust drug with universal applicability and high efficacy, that can be scaled rapidly to high levels of production in order to fight this virus family. Considering the fact that trials and approval process of a novel designed small-molecules take approximately 10 to 15 years in the US (4), it would be a wise choice to repurpose the already well characterized small-molecules to respond to this emergency.

To this date, there is a significant amount of work done to test previously approved antiviral drugs, resulting in a few successes, like Remdesivir by Gilead and Molnupiravir by Merck with modest levels of improvement in treatment outcomes. However, there is still a need for a more comprehensive treatment of SARS-CoV-2 to further decrease the fatality and minimize the spread of virus.

As an early therapeutic strategy, there are already numbers of studies to identify small molecules to inhibit SARS-CoV-2 activity in the host cells. To our knowledge, there are six main strategies/routes to target the viral proteins (5) (Figure 1A): i. Inhibiting the interaction between the viral spike (S) protein and angiotensin-converting enzyme 2 (ACE2) receptor of the host (6, 7). The initial attachment of the virus to the host cell is initiated by this interaction (8). ii. Inhibiting S protein cleavage by proteases. Following receptor binding, the virus gains access to the host cell cytosol by acid-dependent proteolytic cleavage of S protein by TMPRSS2, trypsin, cathepsin L (catL) or other proteases at the S1/S2 boundary and S2’ (8). iii. Blocking the entry of the virus into the cytosol by inhibiting the fusion of virus to the endosomal membrane. Following proteolytic cleavage, the heptad repeat 1 (HR1) & heptad repeat 2 (HR2) regions of the S2 domain interact to form a six helical bundle which plays an important role in the fusion to the membrane. HR1 & HR2 regions can be targeted to prevent viral entry into the host cell (9). iv. Inhibiting the proteases essential for proteolytic processing of viral polyproteins (10, 11). After entering the host cell, the virus continues its lifecycle by translating its genes which express two co-terminal polyproteins, pp1a and pp1ab. Proteolytic processing of viral polyproteins by main protease (Mpro) and papain-like protease (PLpro) results in nonstructural proteins which basically inhibit most of the function of the host cells. v. Inhibiting reverse transcription of the viral genomic material and its multiplication (12). Finally, vi. Inhibiting assembly, packaging and release of newly formed viruses (12).

**Figure 1.**
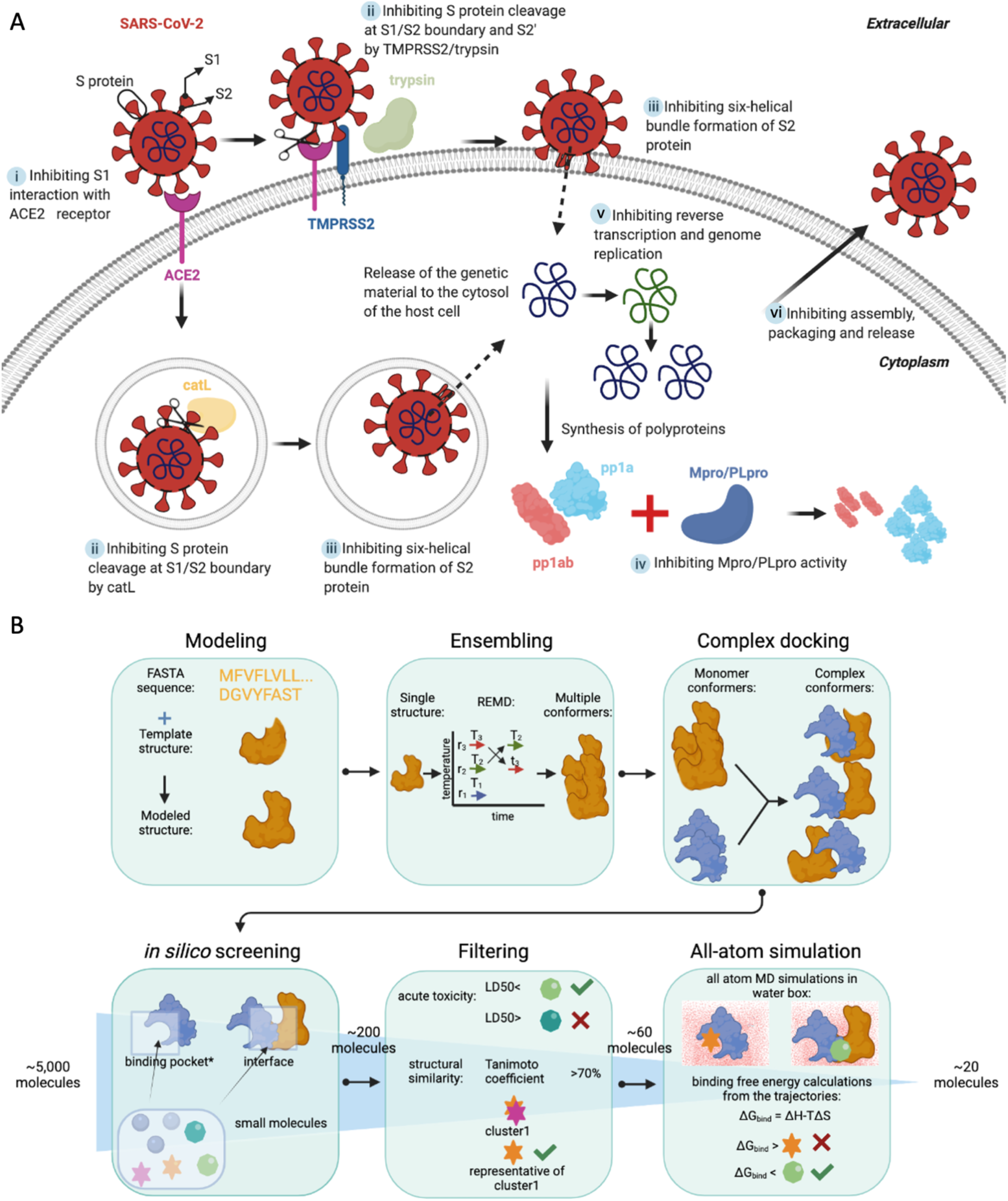
The targeting strategy and pipeline. (A) A summary of viral protein targeting strategies. Among six main strategies denoted here as i, ii, iii, iv, v and vi, we focused on ii. (B) The computational pipeline flow used for screening drugs starting from the target fasta sequence to the bench. r in the ensembling panel stands for replica, while T in the same panel stands for temperature. * binding pockets of the monomers correspond to the interface sites of each binding partner in the complex The number of small molecules at the beginning of the *in silico* screening and in-between each step is written before and after the steps. *in silico* screening starts with 5,000 molecules, this number decreases to 200 after the screening. Then, filtering narrow this number down to 60 molecules. MD simulations and binding free energy calculations results in 20 molecules to be evaluated on the bench. This figure is created with BioRender.

Here, we focused on inhibiting viral entry (route ii, Figure 1A) by performing a structure-based screening of FDA approved drug compounds against the proteases trypsin, catL, and TMPRSS2 as well as the viral S protein. S protein consists of two main domains; S1 domain responsible for receptor binding and the S2 domain mediating membrane fusion. In the case of SARS-CoV, the S protein is proteolytically cleaved at the S1/S2 boundary (13). However, recent studies have revealed that cleavage at a secondary site (S2’) is also critical for viral activation by serine proteases, like TMPRRS2 and trypsin, and entry into the host cell (13, 14). Therefore, we aim to identify small molecules to be repurposed to disrupt S protein cleavage at both S1/S2 boundary and S2’. Even though the SARS-CoV endosomal entry requires catL protease (15), it has been shown that TMPRRS2-mediated activation, governed by the S2’ site, is required for viral pathogenesis (16, 17). Our approach was to target either the S protein and/or proteases to inhibit both endosomal (catL) and TMPRRS2-mediated entry of the virus. Recent efforts have already produced all-atom structures of the novel SARS-CoV-2 S protein, showing a well-established structural region at S2’, which can be deployed to perform molecular drug screening. Furthermore, this secondary site is conserved between the SARS variants (unlike the primary site), and therefore allows for broader applicability against other strains of the virus.

In this study, we obtained structural ensembles of protease-S protein complexes that capture the inherent configurational diversity using replica exchange molecular dynamics (REMD) simulations, and used those structures for *in silico* small molecule screening. This approach allows us explore thermally accessible configurations other than just the lowest energy structure, which are likely to play important roles in molecular recognition (18). In addition to targeting the S protein from the original strain, we also included a recently observed S protein mutation to our analyses. This mutation is identified as a result of analysis of the HR1 mutation occurrences in 34,805 the SARS-CoV-2 genomic sequences in GISAID (19). The study revealed a dramatic increase for the D936Y mutation, which was particularly widespread in Sweden and Wales/England. We spotted this mutation as close to the trypsin binding site of S2’, and therefore we obtained ensembles of configurations for trypsin-mutant S protein complex.

We used the structural ensemble of the proteins to screen approximately 5,000 small molecule structures. These structures were curated from various drug libraries and they were all FDA approved small molecular drugs. However, the recurring structures occurring more than one database were not removed and different confirmations of the same small molecular drugs were included in the screening. From this ensemble docking procedure, we ranked the drug molecules based on predicted binding affinities to structures of unbound proteases and S proteins, as well as protease-S protein complexes, and we identified the top ∼200 molecules with < −7.4 kcal/mol binding free energies.

Furthermore, we filtered down the list to 55 molecules by clustering based on chemical, structural and functional similarity, and eliminating molecules with high toxicities. The top 55 molecules were subject to all-atom molecular dynamics (MD) simulations to assess the stability of the interaction in explicit water, and for stable complexes binding free energies were estimated using the Molecular Mechanics Poisson–Boltzmann Surface Area (MM-PBSA) method to produce a list of lead hits. Finally, we tested our lead candidates experimentally, at least in one case (against catL), for their potential to inhibit protease activity. Indeed, the experiments revealed catL inhibition for two of the five molecules that were predicted in silico, highlighting the potential of this pipeline (Figure 1B) for high throughput drug screening.

## 2 Results

To perform docking, we generated an ensemble of receptor configurations using REMD simulations. To that end, we performed root-mean-square deviation (RMSD) clustering of the structures obtained from the simulations, and chose the central structures from the top three clusters, which accounted for more than 70% of all conformational states (Figure 2 and Figure 3). Over 205,000 docking calculations were then performed to dock 5,000 small molecules to our ensembles of configurations for our 17 monomer and 24 complex structures. The top 20 hits for each structure were selected. Below we discussed specific results of docking calculations for monomer and complex structures, separately.

**Figure 2.**
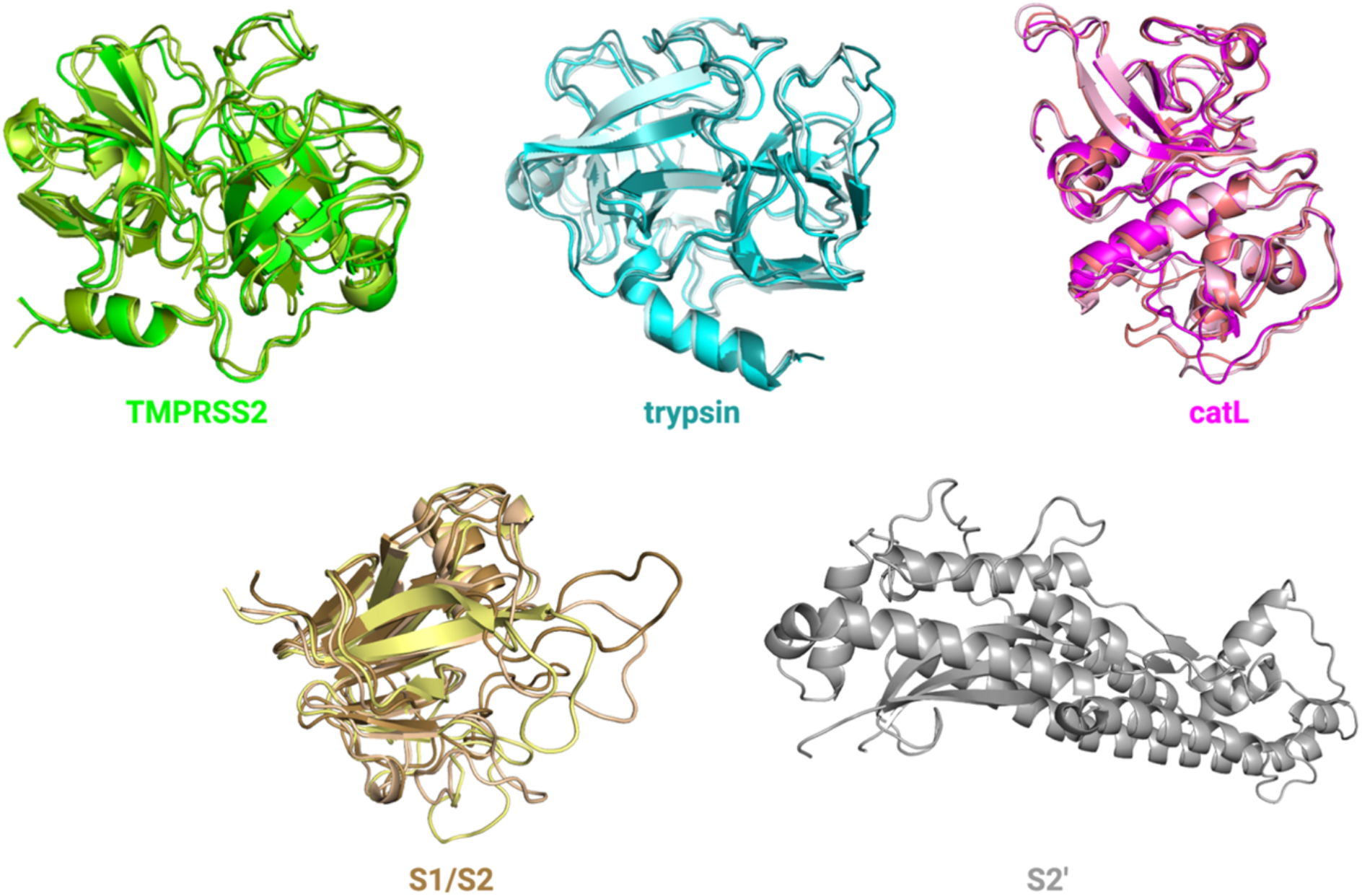
Ensembles of configurations of unbound proteases and S protein cleavage sites. Three configurations for TMPRSS2, trypsin, catL and S1/S2 as well as one configuration for S2’ were used for *in silico* screening.

**Figure 3.**
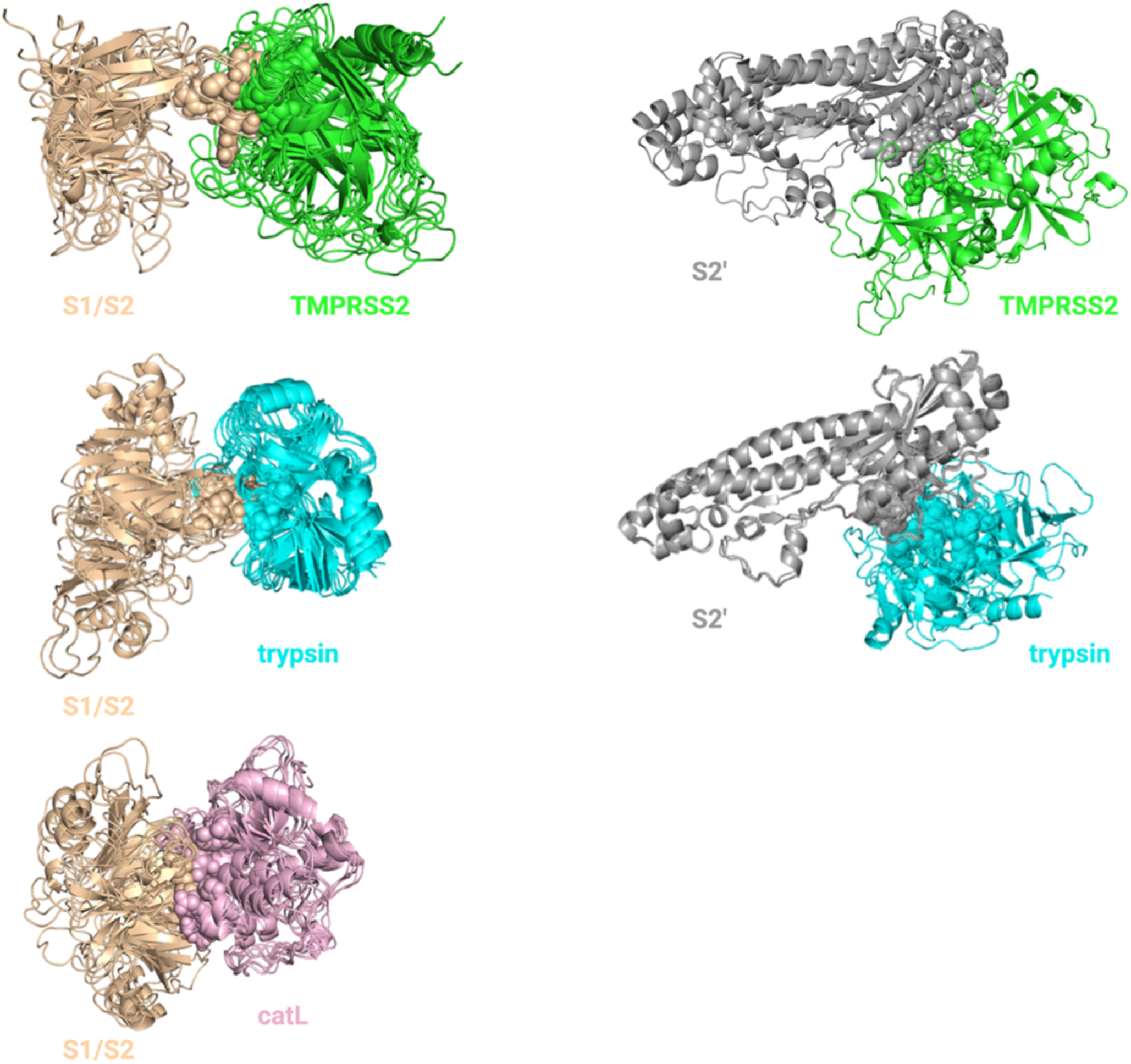
Ensembles of configurations of protease-S protein complexes. Four configurations for TMPRSS2-S1/S2 complex, two configurations for TMPRSS2-S2’ complex, six configurations for trypsin-S1/S2 complex, three configurations for trypsin-S2’ complex, six configurations for catL-S1/S2 complex were used for *in silico* screening. Catalytic sites of the proteases and cleavage sites of the S protein are shown with spheres.

### 2.1 Evaluating the *in silico* docking pipeline incorporating an ensemble of receptor configurations

To evaluate the performance of this method, we took advantage of the Directory of Useful Decoys (DUD) dataset (20), which contains validated list of active and decoy compounds for a number of proteases, including one of our targets (Trypsin). Using the ensemble generated for Trypsin, we were able to dock the active (47) and decoy molecules (1652), and compare the performance against docking to the crystal structure, which we refer to as the naïve method. The results are summarized in terms of a Receiver Operating Curve (ROC) shown in Figure 4. The naïve method performs decently as characterized by an Area Under the Curve (AUC) of 0.71, which is in good agreement with previously published performance of Vina (AUC = 0.72) (21). Remarkably, the ensemble method showed a significant improvement in performance with an AUC of 0.84, aided by molecular simulations in explicit water. Therefore, we proceeded with this ensemble method for screening of the drugs against our protease and spike protein targets. In the future, we plan on validating this method to a wide range of targets from DUD database to comprehensively evaluate its performance.

**Figure 4.**
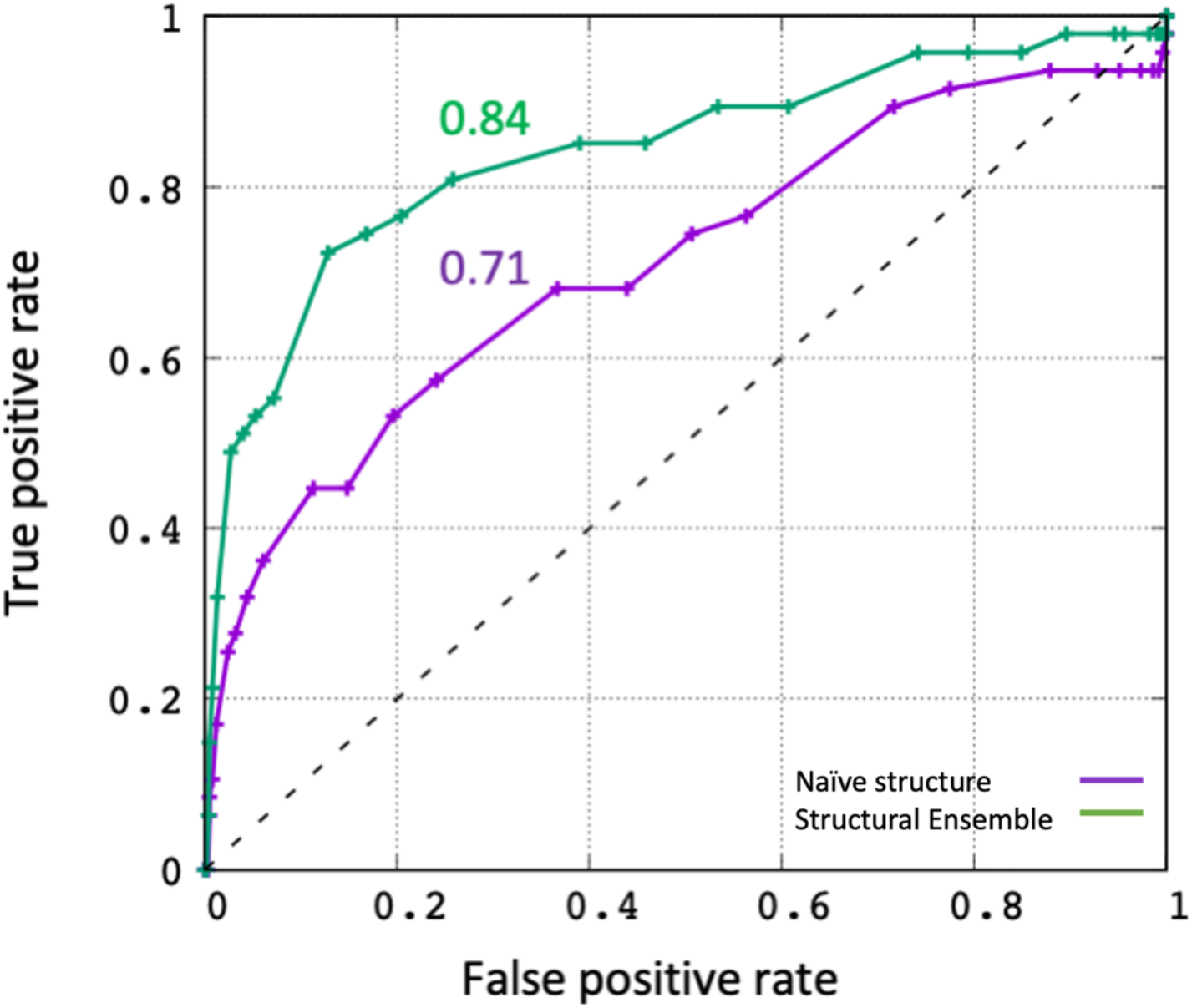
Receiver Operating Curves (ROC) to evaluate performance of the *in silico* screening. Docking of active (47) and decoy (1652) molecules were performed against the crystal structure of Trypsin (purple) and against the structural ensemble generated by REMD simulations (green). The Area Under the Curve (AUC) values are also shown.

### 2.2 Identifying the top molecules to target unbound proteases and S protein, and protease-S protein complexes

The S2’ site of the S protein was not subject to REMD simulations, as the target region is a rigid alpha helix bundle. Therefore, the ensemble of monomer receptors comprised a total of 17 structures are as listed: 3 each of trypsin, catL, TMPRSS2, S1/S2-a, S1/S2-b, and one of S2’ & S2’_D936Y. S1/S2-a refers to docking against S1/S2 boundary cleavage site targeted by TMPRSS2 and trypsin (R685-S686), while S1/S2-b indicates docking against S1/S2 boundary cleavage site targeted by only catL (T696-M697). S2’, by itself, is the secondary cleavage site on S protein targeted by TMPRSS2 and trypsin.

85,000 docking calculations using the 17 conformations of S protein and proteases were performed and the top 19-21 hits for each ligand was listed in Table S1. The docking scores varies from −13.6 to −7.4. From the list, 96 distinct molecules (hereinafter referred to as ligands) were identified targeting either/both S protein cleavage sites or/and catalytic sites of proteases. Considering the similarity between catalytic sites of TMPRSS2 and trypsin, we expected to see some common molecules targeting both of these proteases, as seen in the table.

The total of 24 complex structures are as listed; trypsin_1-S1/S2_1, trypsin_2-S1/S2_1, trypsin_3-S1/S2_1, trypsin_1-S1/S2_2, trypsin_2-S1/S2_2, trypsin_3-S1/S2_2, trypsin_1-S2’_1, trypsin_2- S2’_1, trypsin_3- S2’_1, trypsin_1- S2’_D936Y_1, trypsin_2- S2’_D936Y_1, trypsin_3- S2’_D936Y_1, catL_1- S1/S2’_1, catL_2- S1/S2’_1, catL_3- S1/S2’_1, catL_1- S1/S2’_2, catL_2- S1/S2’_2, catL_3- S1/S2’_2, TMPRSS2_1- S1/S2_1, TMPRSS2_2- S1/S2_1, TMPRSS2_1-S1/S2_2, TMPRSS2_2- S1/S2_2, TMPRSS2_1- S2’, TMPRSS2_2- S2’. Here is an example to clarify the nomenclature, trypsin_1-S1/S2_1 refers to complex between the first configuration of trypsin and the first configuration of S1/S2 boundary domain of S protein.

120,000 docking calculations using these 24 complexes were performed and the top 19-21 hits for each ligand was listed in Table S2. The docking scores varies from −13.7 to −9.2. From the list, 151 distinct ligands were identified targeting S protein-protease interaction interfaces.

### 2.3 Curating a more refined list of ligands according to their toxicity, structural and functional similarity

From the 96 and 151 distinct ligands from in silico screening results for the monomer and the complex structures, respectively, we used three different selection criteria to arrive at a more refined list of ligands. The first selection criterion is based on the toxicity of the ligands. As mentioned before, FDA approved drugs were used in this study to meet the urgent need for a treatment. To further pursue this approach, the ligands with high acute toxicity rate (such as chemotherapeutics), which cannot be delivered at high concentrations, were given the least preference. The second and third selection criteria are based on the structural and functional similarities of the ligands, respectively. Chemical clustering of ligands was performed according to their Tanimoto coefficients with a similarity cut-off of 70%. Furthermore, the mechanism of action was assigned to each ligand. For example, Tyrosine kinase inhibitors, NS5A inhibitors, ergosterol binders were among the most commonly occurring action mechanism of the ligands. Finally, the least toxic ligands with a unique mechanism of actions were selected from each cluster. The atopical ligands are eliminated. The refined list of 55 ligands with their LD50s and functions is shown in Table 1. Interestingly, drugs previously proposed or/and used for COVID-19 treatment such as Remdesivir, Gleevec, Saquinavir, Digoxin, Camostat, Drospirenone, Dexamethasone M and Dihydroergotamine are among these 55 ligands. However, their potential for blocking viral entry needs to be further explored.

**Table 1.**
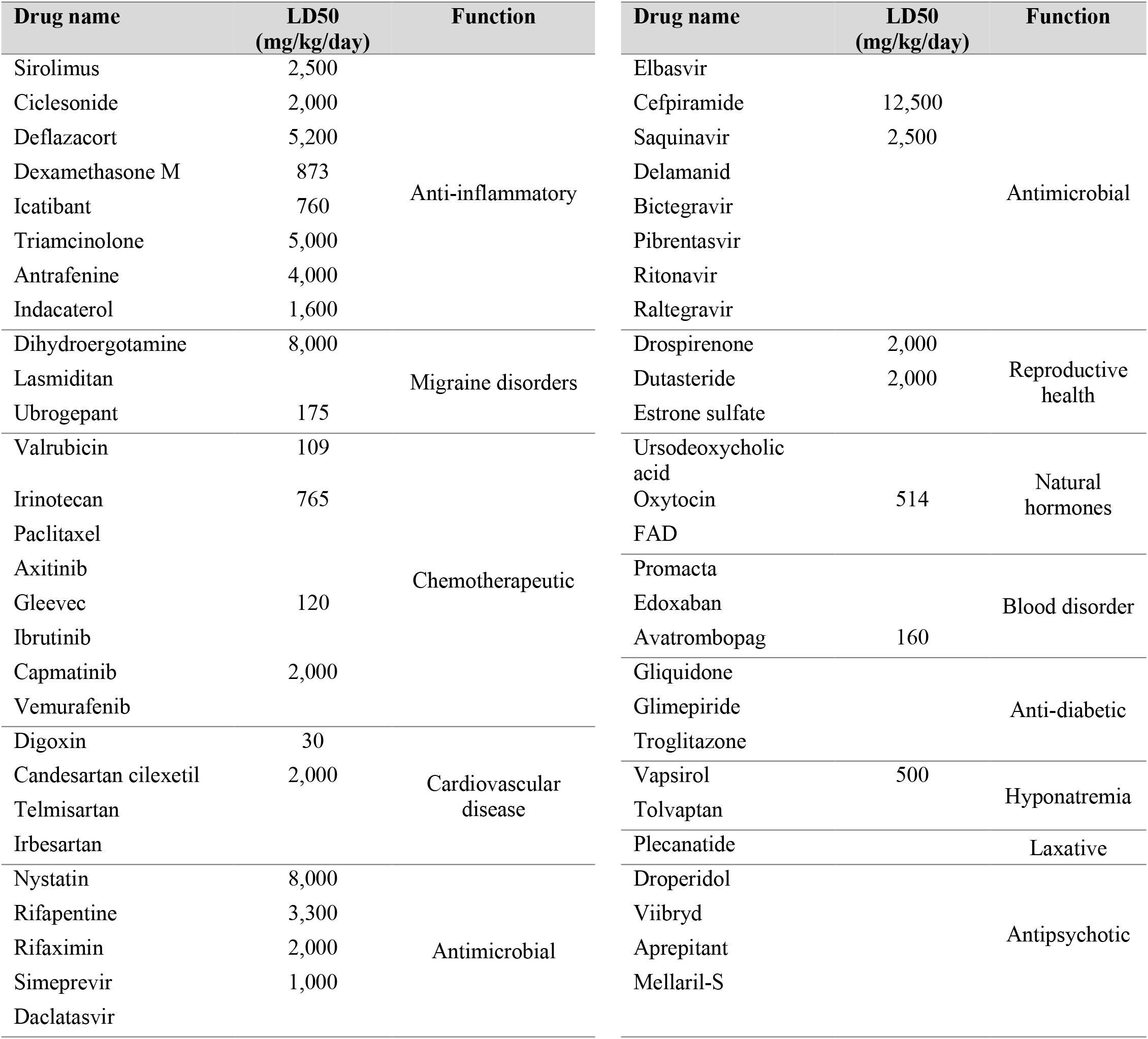
The list of ligands refined according to their toxicity, structural and functional similarity

### 2.4 Identifying the best binding ligands to inhibit proteolytic cleavage of S protein

Considering both TMPRSS2 and trypsin are serine proteases, some of the 55 ligands were targeting more than one complex or monomer. There were 173 initial configurations consisting of ligand-monomer or ligand-protease-S protein complexes. Atomistic MD simulations were performed on these complexes for 100 ns. Using a MMPBSA energy cut-off (ΔG < 0) (Table S3), we identified a small number of ligands for each protease and S protein to specifically inhibit proteolytic cleavage of S protein (Table 2 and Figure 5). The ligands listed in Table 2 can be seen in Figure 5, which shows the positioning of the ligands in the interfaces of the complex or cleavage/catalytic sites of S protein/proteases.

**Table 2.**
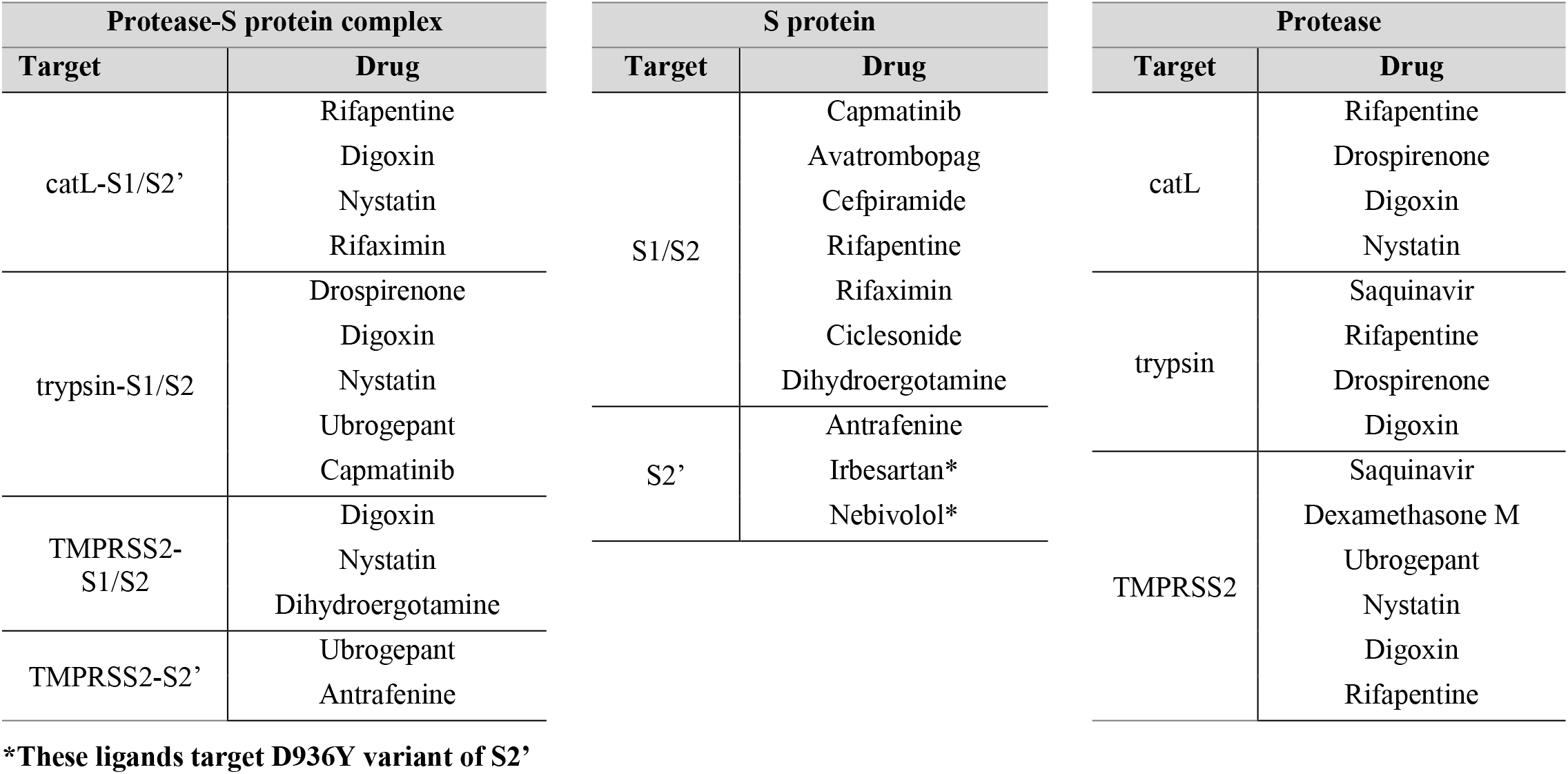
The ligands and their targets obtained by MD simulations and Δ***G*** calculations

**Figure 5.**
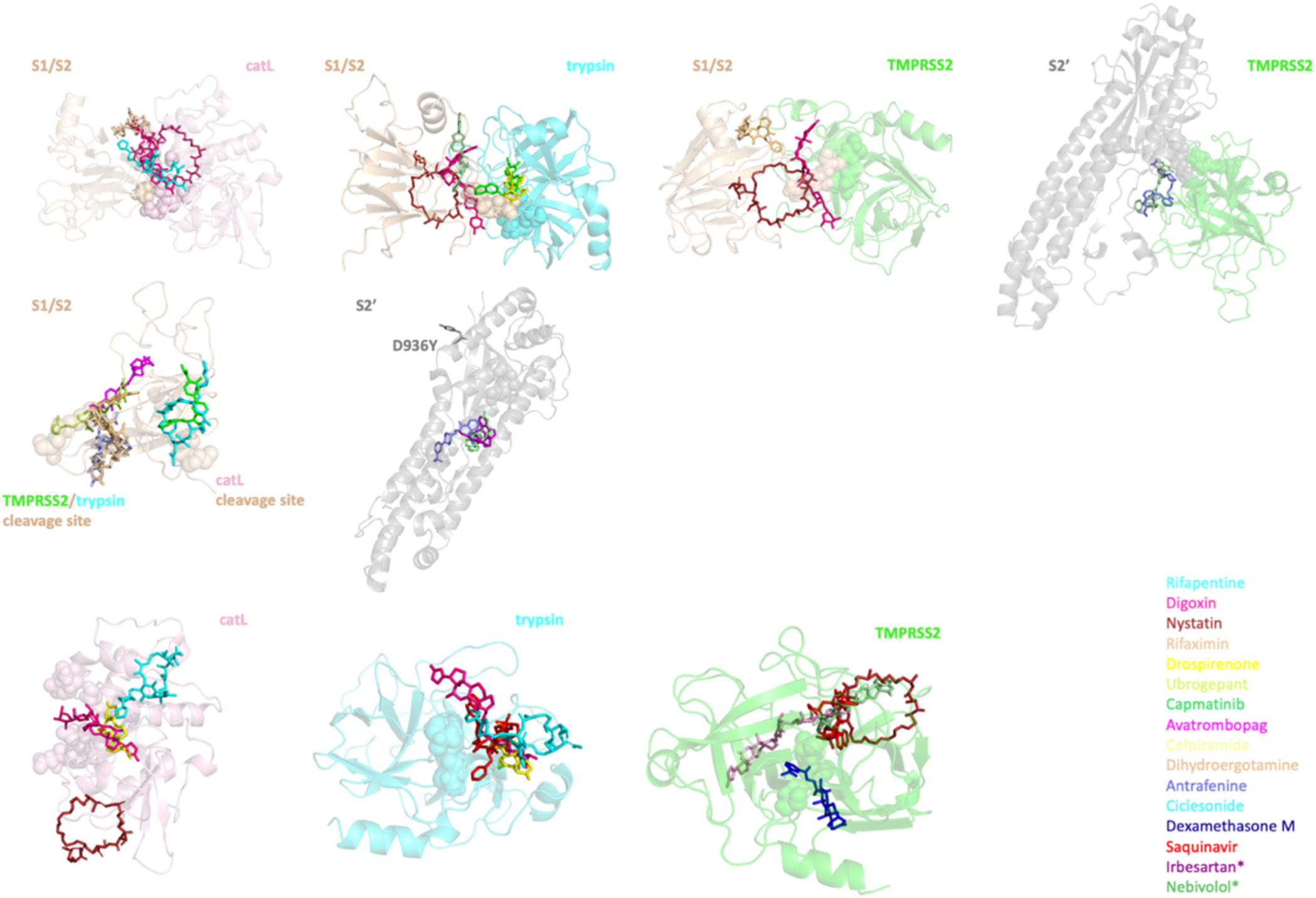
The ligands in complex with their targets. The ligands bound to their targets are shown in stick representation. Their color-coded list can be seen on the right. Catalytic sites of the proteases and cleavage sites of the S protein are shown with spheres. Two different cleavage sites on S1/S2 region of S protein are labeled on the figure. *Irbesartan and Nebivolol are specific to D936Y variant of S protein

The *in silico* screening approach has allowed us to narrow down the possible viral inhibitors from ∼5,000 to ∼ 20, which can be tested experimentally for their efficacy to block viral entry into host cells. To demonstrate proof of concept, we proceeded to validate some of our lead hits experimentally at least in one case. We designed an in vitro protease activity assay for catL, and observed the possible inhibitory effects of the four drug molecules that are predicted to interact with the active site of the proteases. We introduced the four putative drugs and a known inhibitor (as positive control) over a range of drug concentrations. As expected, the known inhibitors had a significant decrease in fluorescence activity. Excitingly, we found that Nystatin and Rifapentine had measurable effects on the activity of catL. Specifically, Nystatin has an IC50 of ∼2 μM and Rifapentine had an IC50 of ∼30 μM for inhibiting the protease activity (Figure 6A). In our handful of lead hits, we found at least two molecules that show potential to inhibit protease activity in μM concentrations. These results are also corroborated by the MD simulations, as evidenced by the RMSD plots of the drugs bound to catL (Figure 6B). Interestingly, Nystatin exhibits most stability bound to the protease indicated by the lowest RMSD for majority of the simulation. While we observed configuration rearrangement toward the end of the simulation, it still remained in close proximity to the active site (the catalytic cysteine is shown in yellow). While Rifapentine, shows a large jump in the RMSD from the initial configuration, it is interesting to note that in the higher RMSD state, the molecule resides close to the active site, and flips out of the active site while still being bound to the protease. This behavior offers a potential molecular explanation for partial/weaker inhibition of the protease by Rifapentine.

**Figure 6.**
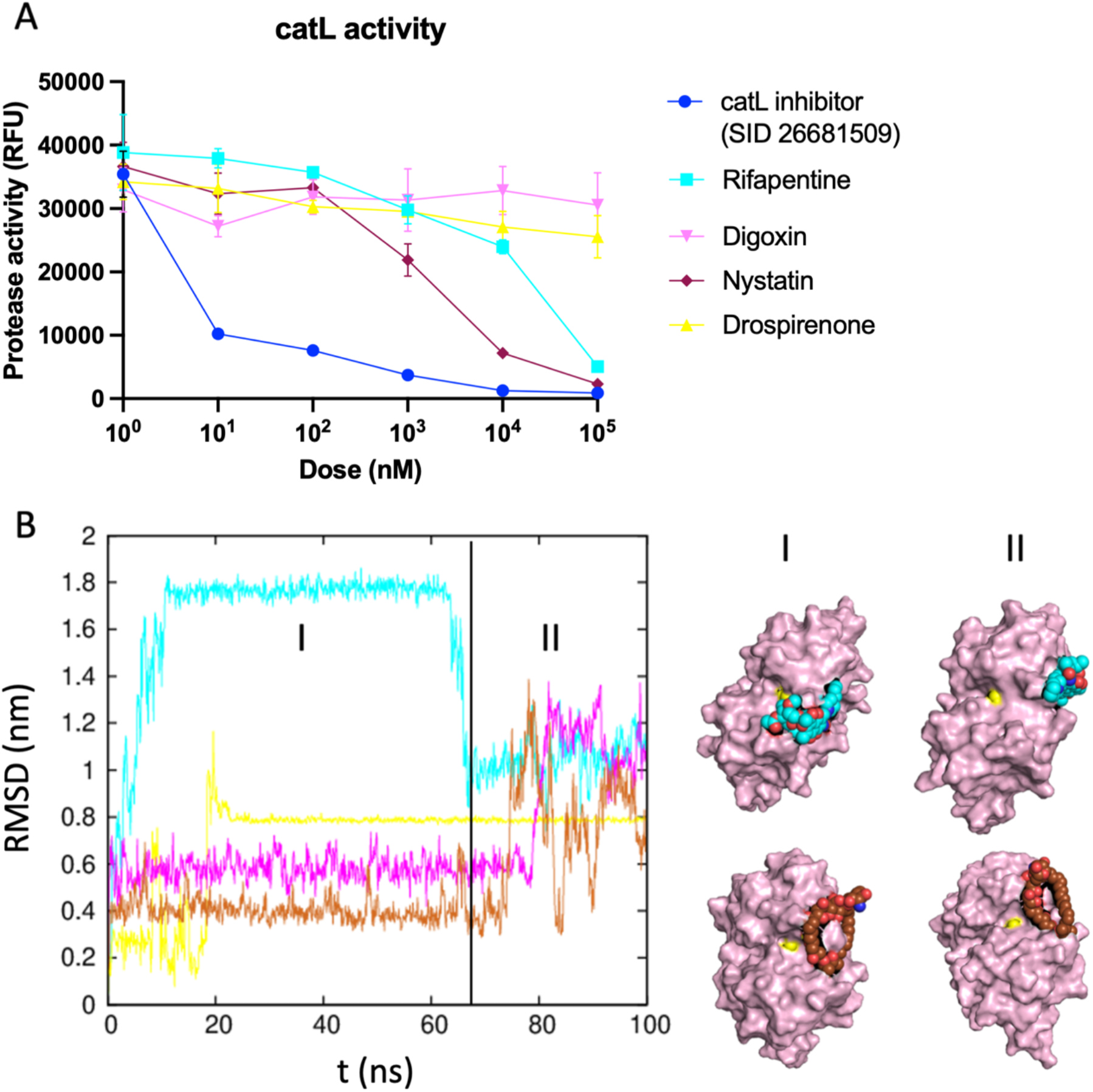
Changes in protease activity of catL upon introducing our proposed small molecules and mechanistic explanation of their effects. (A) catL protease activity upon addition of SID 26681509, Rifapentine, Digoxin, Nystatin and Drospirenone. RFU stands for relative fluorescence units. (B) RMSD trends of the complexes between catL and tested drugs (Rifapentine, Digoxin, Nystatin and Drospirenone). On the right, two snapshots of our lead drug candidates, Rifapentine (cyan) and Nystatin (brown), in complex with catL (pink) from the start (I) and the end (II) of the simulations are shown. It suggests that interaction of Rifapentine and Nystatin to the catL active site (shown with yellow sphere) stabilizes the whole complex.

## 3 Discussion

While the vaccines have provided adequate immunity against the virus, new variants, like Omicron with several mutations in the spike protein can be handled comprehensively by establishing a therapeutic strategy that will remain effective against future variants. TMPRSS2, trypsin and catL are considered as potential targets to inhibit SARS-CoV infection dating back to 2003 (22, 23). Inhibition of proteolytic cleavage of S protein of SARS-CoV-2 by those proteases emerges as crucial since 1) it prevents the viral genomic material release to the host cell cytoplasm. 2) it is effective for both extracellular and endosomal viral entry.

In this study, we used homology modeling and REMD to generate an ensemble of configurations for ensemble docking. This approach enabled us to sample a wide variety of conformations of the target proteins (S protein and proteases). The structures chosen for docking following RMSD clustering accounted for more than 70% of the conformational states sampled, which produced the diversity needed for ensemble docking. We screened 5,000 small molecules against our ensembles of configurations for our 17 monomer and 24 complex structures, summing up to over 205,000 docking calculations. Following docking, we narrowed down the lead hits by filtering them based on acute toxicity (LD50) and Tanimoto coefficient of our ligands to identify unique ligands with low toxicity, leading us to a refined list of 55 drugs. This narrowing down allowed to study the 55 drugs in complex with their targets (173 complexes) in more detail using all-atom explicit water MD simulations to assess the stability of the interaction and quantify the binding energy in the presence of the solvent explicitly. The RMSD plots and the binding energies from the MD simulations were used to finally rank the drug-target interactions, which allows further experimental investigation of a short list of drugs for their antiviral potential.

As we mentioned SARS-CoV-2 variants can acquire mutations that increase their virulence, like enhanced binding to the ACE2 receptor. Therefore, if the treatment regimen involves targeting the ACE2 binding domain of the S protein, its efficacy might decrease when the mutations alter the binding to the drug. In this study, we have targeted the human proteases and cleavage sites of S protein, which tend to be conserved even in the variants. Nevertheless, we identified one variant with a mutation D936Y on S protein, which is close to the S2’ cleavage site. We modeled the variant with the mutation, and identified Irbesartan and Nebivolol as potential drugs specific to D936Y variant.

The computational pipeline allowed us to screen a large number of drugs, and narrowed the list of interest to a few that can be tested at the bench. Two of the four drugs, Nystatin and Rifapentine, showed measurable inhibitory effects on catL activity. Nystatin is an antifungal that is usually used to treat fungal infection in mouth, which rifapentine is an antibiotic used for treating tuberculosis. It is important to note that these drugs have not been tested for their potential for treat COVID19, and the authors recognize such premature applications of the drug can be harmful and result in shortage of the drugs for patients who need them currently. We are hoping that extensive testing of these drugs for their potential to treat COVID19 is conducted together with the other recently proposed potential proteolytic inhibitory molecules (24, 25), starting from in vitro studies for their potential to block viral entry.

In this study, we generated a computational pipeline for screening drugs from the target fasta sequence to the bench (Figure 1B). Combination of the computational and experimental techniques enabled us to screen all the FDA approved drugs and identify some drugs that can effectively target our proteins of interest. Our data recommends a deeper investigation of around a dozen drugs for their potential to inhibit both endosomal (catL)-mediated entry of the virus. Many effective vaccines are being applied worldwide and they significantly reduce that number of new cases in countries with rapid vaccination rates. However, a treatment is still in need since the vaccine protection is not 100% and there is inequitable vaccine distribution across the world. Therefore, we believe our proposed regimen will help to meet this need.

## 4 Material and Methods

### 4.1 Computational methods

#### 4.1.1 Initial configurations of SARS-CoV-2 S protein, TMPRSS2, trypsin and catL

The crystal structure of SARS-CoV-2 spike ectodomain has been already solved (PDB ID: 6vyb (26)), however it has some unstructured missing region. Therefore, the structure of whole length S protein was modeled using SWISS-MODEL web server (27); the crystal structure, 6vyb, and FASTA sequence (NCBI Reference Sequence: YP_009724390.1) of SARS-CoV-2 S protein were used as template. S1/S2 boundary and S2’ structures were adapted from our whole length S protein model. S1/S2 boundary structure covered residues from 293 to 326 and from 591 to 700, S2’ structure consisted of residues from 715 to 1070 (Figure 2, bottom panel).

TMPRSS2 extracellular domain was modeled giving Hepsin crystal structure (PDB ID: 1z8g (28)), a closely-related serine protease with more than 50% sequence identity, and FASTA sequence of TMPRSS2 (UniProt ID: O15393) as template using SWISS-MODEL.

catL and trypsin crystal structures were already available in PDB (PDB ID: 2xu1 (29) and 1h4w (30), respectively) and they were used in this study.

#### 4.1.2 Replica exchange MD simulations

REMD simulations were performed to generate an ensemble of configurations for subsequent protein-protein docking. Simulations were performed using GROningen MAchine for Computer Simulations (GROMACS-2018) on an Exacloud cluster at Oregon Health & Science University (Portland, OR).

The protein structures were centered in a 3D periodic cubic box with initial dimensions ∼ 7-8 nm in the three orthogonal directions. The box was then solvated with ∼ 12,000 to 14,000 water molecules, depending on the size of the box, and a few Na^+^ or Cl^-^ counterions, depending on the charge of the protein, to keep the system charge neutral. After an initial equilibration of 1 ns, a REMD schedule was setup with 60 windows in the temperature range of 300-425K. The temperature schedule was generated using the webserver (http://virtualchemistry.org/remd-temperature-generator/). The REMD simulations were then performed on the 60 windows for 15 ns with exchanges between neighboring windows tried every 0.4 ps, and the protein co-ordinates stored every 50 ps. The first 5 ns were ignored, and the last 10 ns, consisting of 200 frames were used for clustering analysis.

Top three configurations from each ensemble covering more than 70% of all configurations were used (except for S2’) for *in silico* screening of three unbound protease structures and one S1/S2 boundary structure. RMSD values within each ensemble vary from 0.9 to 1.3 Å. For the screening of S2’, one equilibrated configuration was used (Figure 2)

#### 4.1.3 Ensemble docking for S protein–protease complex modeling

The configurations generated by REMD for two cleavage domains of S protein (S1/S2 and S2’), TMPRSS2, trypsin and catL were used to obtain ensembles of configurations for each S protein-protease complex. Top two configurations from S1/S2 boundary and TMPRSS2 ensembles and top three configurations from trypsin and catL ensembles covering ∼70% of all configurations were selected for the docking. RMSD values within each ensemble vary from 1.9 to 5.9 Å. One equilibrated S2’ configuration was used. TMPRSS2 and trypsin were docked to both S1/S2 boundary (R685-S686) and S2’, while catL was only docked to a different cleavage site on S1/S2 boundary (T696-M697) using HADDOCK web server (31) (Figure 3). Active residues for flexible docking were defined as follows; catalytic triad of TMPRSS2 and trypsin, active sites of catL identified in Fujishima et al (32), and cleavage sites on S1-S2 boundary and S2’. The cleavage sites on S1/S2 boundary and S2’ for TMPRSS2, trypsin and catL were obtained by aligning protein sequences of SARS-CoV S protein (UniProt ID: P59594) and SARS-CoV-2 S protein. The sequences of previously defined cleavage sites on SARS-CoV (14) were conserved in SARS-CoV-2 but residue numbers were different (Table 3). In order to obtain, mutant S2’ monomer and mutant S2’-trypsin complexes, PyMol (version 2.3.2) built-in Mutagenesis tool was used.

**Table 3.** Cleavage sites on SARS-CoV and SARS-CoV-2 S proteins

| SARS-CoV S protein | SARS-CoV2 S protein | Cleaved by |
| --- | --- | --- |
| R667-S668 | R685-S686 (S1/S2 boundary) | TMPRSS2 & trypsin |
| R797-S798 | R815-S816 (S2') | TMPRSS2 & trypsin |
| T678-M679 | T696-M697 (S1/S2' boundary) | catL |

#### 4.1.4 Small molecule libraries and structure based *in silico* screening

Clinically approved small molecules from the following drug libraries were used for screening; ZINC library of ∼1,650 molecules (33), DrugBank of ∼ 2,500 molecules (34), The Binding Database of ∼1,340 molecules (35) and ChemBridge of ∼50 molecules (top 50 hits from Elshabrawy et al (15)) obtained from Chembridge Corporation (San Diego,CA). All SDF files from each library were converted to PDBQT using Open Babel software (36).

AutoDock Vina (version 1.1.2, Linux) (37) was used for the docking of small molecules to our configurations. Two approaches were followed for the docking process. First, total of 17 configurations of unbound proteins (16 WT S protein and protease monomers, and 1 mutant S protein monomer) from REMD were used for *in silico* screening to study the binding of small molecules to the unbound receptors. Moreover, 24 S protein−protease complexes (21 WT S protein-protease complexes and 3 mutant S protein-proteases complexes) obtained by flexible docking were used for the screening. These two approaches allowed for *in silico* screening against each PPI partner individually at the interface site, as well as opportunity to evaluate the possibility of the small molecules destabilizing S protein-proteases complexes. Hydrogens were added to all receptors and grid box dimensions were defined according to the size of receptor binding pockets or interfaces.

#### 4.1.5 Filtering

Filtering of our ∼160 lead candidates were done using chemical similarity & toxicity. Chemical similarity was quantified using the Tanimoto coefficient as calculated using the Molecular Operating Environment (MOE). The acute toxicity as quantified by the mice lethal dose (LD50) was downloaded from FDA drug reports & go.drugbank.com. The mechanism of action data was curated from PubChem database (38). The least toxic drugs were selected from clusters that were generated using a 70% chemical similarity, yielding a list of ∼ 55 molecules.

#### 4.1.6 Force field generation

All-atom AMBER-type force-field parameters were generated for the lead 55 molecules. We created a pipeline to start from a pdb file of the lead molecules to producing gromacs type forcefields for the protein-drug complexes. This pipeline included mostly AMBER tools (39), to calculate partial charges at HF/6-31G (AM1-BCC) (40), and use the General AMBER Force-Field (GAFF) (41) for bonded and van der Waals interactions, after ensuring the right protonation state for the molecules.

#### 4.1.7 All-atom MD simulations

A total of 17.0 μs simulations, each with 100 ns, were performed using the GROningen MAchine for Computer Simulations (GROMACS-2018) on an exacloud cluster at Oregon Health & Science University (Portland, OR). MD simulations in an aqueous environment were carried out for a total of 173 initial configurations of small molecule-protein complexes. The TIP3P water model was used to simulate the system in an aqueous environment with proper number of counterions (Na+ or Cl-) to ensure charge neutrality. A 3D periodic box was used to center the complex with at least 1.0 nm from the edge. In the equilibration and production runs, the temperature was maintained at 300K and the pressure was maintained at 1 bar, using V-rescale thermostat (42) and Parrinello-Rahman barostat, respectively. The MD simulations incorporated leap-frog algorithm with a 2 fs time-step to integrate the equations of motion. The long-ranged electrostatic interactions were calculated using particle mesh Ewald (PME) (43) algorithm with a real space cut-off of 1.2 nm. LJ interactions were also truncated at 1.2 nm. To monitor the systems to reach equilibration, root-mean-square deviation (RMSD) of the complex structures were calculated as a function of time. Furthermore, to analyze the interaction of the drug with the proteins, we performed RMSD of the drug during the course of the simulation, while minimizing the RMSD of the protein monomer/complex.

#### 4.1.8 MM-PBSA calculations

The average binding free energy is derived from the Gibbs free energy,

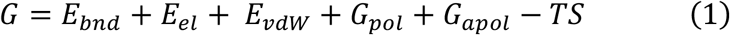

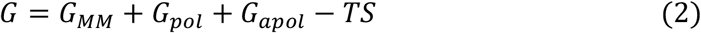

where G_MM_, G_pol_ and G_apol_ denote the Molecular Mechanics, polar and apolar contributions, respectively (44). TS, the entropic contribution, was not included in our calculations. GROMACS g_mmpbsa package was used to obtain the G_MM_, G_pol_ and G_apol_ contributions (45, 46). The package allows user to define the protein and ligand during the calculation and measure the average binding free energy between the ligand and protein as follows;

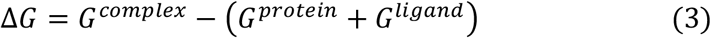

### 4.2 Experimental methods

#### 4.2.1 catL Activity Assay

rh catL was diluted to 40 µg/mL in Assay Buffer (50mM MES, 1mM EDTA, 0.005% (w/v) Brij-35, pH 6.0). Diluted rh catL was incubated on ice for 15 minutes. Incubated 40 µg/mL rh catL was diluted to 40ng/mL in Assay Buffer. 25uL of catL inhibitor (SID 26681509) was loaded in 96 black well transparent bottom plate, then 25*μ*L of cathepsin L (1:1 dilution) was loaded and incubated at 37°C for 30 minutes. The substrate was diluted to 80 µM in Assay Buffer, then 50*μ*L of substrate was added to each well and incubated as wrapped in foil at 37°C for 1 hour. Three replicates for each drug concentration were used. They were imaged on TECAN. All tested drugs were diluted in DMSO to 2mM (5*μ*L into 50*μ*L) and then added 5*μ*L to each positive control well.

## 6 Author Contributions

Conceptualization; ESO, SVR, data curation; ESO, formal analysis; ESO, HHL, AY, SVR, funding acquisition; ESO, SVR, methodology; ESO, HHL, AY, SVR, writing; ESO, AY, SVR

## 8 Acknowledgments

All simulations had been performed using the high-performance computational facilities an Exacloud cluster at Oregon Health & Science University (Portland, OR). We thank Stefanie Linch for helpful discussions. We thank Dr. Jared M. Fischer for helpful discussions on experimental procedure.

