## Supplementary Data for "*In silico* screening and testing of FDA approved small molecules to block SARS-CoV-2 entry to the host cell by inhibiting Spike protein cleavage"

Supplementary Material

### 1 Supplementary tables

**Table S1.** Top ~20 small molecule hits targeting the monomers obtained by *in silico* screening *

| **trypsin_1** | | | **trypsin_2** | | | | | | **trypsin_3** | | | | | | | |
| --- | --- | --- | --- | --- | --- | --- | --- | --- | --- | --- | --- | --- | --- | --- | --- | --- |
| **Energy** | **Database** | **Name** | | **Energy** | **Database** | | | **Name** | | **Energy** | | | **Database** | | | **Name** |
| -12.1 | dbfda-world_494 | Nystatin | | -12.6 | DrugBank_approved_980 | | | Candicidin | | -13.6 | | | DrugBank_approved_980 | | | Candicidin |
| -10.5 | DrugBank_approved_980 | Candicidin | | -11.3 | DrugBank_approved_1814 | | | Tannic acid | | -11.9 | | | DrugBank_approved_2005 | | | Venetoclax |
| -10.1 | DrugBank_approved_1814 | Tannic acid | | -10.8 | dbfda-world_1573 | | | Ecamsule | | -11.2 | | | dbfda-world_531 | | | Amphotericin B |
| -9.9 | DrugBank_approved_1137 | Quinupristin | | -10.7 | DrugBank_approved_2005 | | | Venetoclax | | -11 | | | dbfda-world_495 | | | Nystatin |
| -9.9 | dbfda-world_1408 | Eribulin | | -10.6 | dbfda-world_1138 | | | Solifenacin | | -11 | | | dbfda-world_256 | | | Digoxin |
| -9.8 | dbfda-world_531 | Amphotericin B | | -10.5 | dbfda-world_256 | | | Digoxin | | -10.9 | | | TheBindingDB_780 | | | Juxtapid |
| -9.8 | DrugBank_approved_2005 | Venetoclax | | -10.5 | dbfda-world_1519 | | | Trypan blue | | -10.6 | | | DrugBank_approved_506 | | | Nystatin |
| -9.8 | dbfda-world_256 | Digoxin | | -10.2 | DrugBank_approved_1597 | | | Canagliflozin | | -10.5 | | | dbfda-world_496 | | | Nystatin |
| -9.7 | DrugBank_approved_2079 | Entrectinib | | -10.2 | dbfda-world_1572 | | | Ecamsule | | -10.5 | | | DrugBank_approved_1814 | | | Tannic acid |
| -9.7 | dbfda-world_495 | Nystatin | | -10.2 | dbfda-world_545 | | | Ergotamine | | -10.4 | | | DrugBank_approved_1409 | | | Eltrombopag |
| -9.7 | dbfda-world_496 | Nystatin | | -10.2 | dbfda-world_494 | | | Nystatin | | -10.4 | | | TheBindingDB_1035 | | | D.H.E. 45 |
| -9.7 | dbfda-world_736 | Sirolimus | | -10.1 | dbfda-world_959 | | | Dutasteride | | -10.4 | | | dbfda-world_545 | | | Ergotamine |
| -9.7 | TheBindingDB_332 | Tannic acid | | -10.1 | DrugBank_approved_506 | | | Nystatin | | -10.4 | | | TheBindingDB_559 | | | Epinephrine |
| -9.7 | dbfda-world_1028 | Rifapentine | | -10.1 | dbfda-world_495 | | | Nystatin | | -10.3 | | | dbfda-world_1534 | | | Paritaprevir |
| -9.6 | dbfda-world_252 | Valrubicin | | -10.1 | DrugBank_approved_1232 | | | Rutin | | -10.3 | | | TheBindingDB_1155 | | | Membraneblue |
| -9.6 | dbfda-world_1534 | Paritaprevir | | -9.9 | TheBindingDB_568 | | | Vontrol | | -10.3 | | | ChemBridge_26 | | | 7786009 |
| -9.6 | dbfda-world_176 | Dihydroergotamine | | -9.9 | DrugBank_approved_555 | | | Ergotamine | | -10.3 | | | DrugBank_approved_1565 | | | Lomitapide |
| -9.5 | TheBindingDB_939 | Oxytocin | | -9.4 | DrugBank_approved_1057 | | | Saquinavir | | -10.2 | | | DrugBank_approved_555 | | | Ergotamine |
| -9.5 | DrugBank_approved_506 | Nystatin | |  |  | | |  | | -10.2 | | | TheBindingDB_955 | | | Tasigna |
| -9.5 | dbfda-world_493 | Nystatin | |  |  | | |  | | -9.1 | | | DrugBank_approved_1057 | | | Saquinavir |
| **catL_1** | | | **catL_2** | | | | | | **catL_3** | | | | | | | |
| **Energy** | **Database** | **Name** | | **Energy** | **Database** | | | **Name** | | **Energy** | | | **Database** | | | **Name** |
| -11.5 | dbfda-world_256 | Digoxin | | -10.6 | dbfda-world_256 | | | Digoxin | | -11.4 | | | dbfda-world_256 | | | Digoxin |
| -10.9 | DrugBank_approved_506 | Nystatin | | -10.3 | DrugBank_approved_506 | | | Nystatin | | -11.3 | | | DrugBank_approved_506 | | | Nystatin |
| -10.8 | dbfda-world_495 | Nystatin | | -10 | dbfda-world_493 | | | Nystatin | | -10.2 | | | dbfda-world_176 | | | Dihydroergotamine |
| -10.5 | DrugBank_approved_980 | Candicidin | | -10 | DrugBank_approved_980 | | | Candicidin | | -10.2 | | | dbfda-world_495 | | | Nystatin |
| -10.1 | dbfda-world_496 | Nystatin | | -9.9 | DrugBank_approved_1137 | | | Quinupristin | | -10 | | | DrugBank_approved_555 | | | Ergotamine |
| -10 | dbfda-world_1534 | Paritaprevir | | -9.8 | dbfda-world_176 | | | Dihydroergotamine | | -9.9 | | | dbfda-world_1299 | | | Midostaurin |
| -10 | dbfda-world_730 | Conivaptan | | -9.7 | dbfda-world_496 | | | Nystatin | | -9.8 | | | DrugBank_approved_1137 | | | Quinupristin |
| -9.9 | dbfda-world_176 | Dihydroergotamine | | -9.3 | TheBindingDB_821 | | | Camptosar | | -9.8 | | | dbfda-world_531 | | | Amphotericin B |
| -9.6 | DrugBank_approved_1137 | Quinupristin | | -9.3 | DrugBank_approved_540 | | | Amphotericin B | | -9.7 | | | TheBindingDB_821 | | | Camptosar |
| -9.6 | dbfda-world_1519 | Trypan blue | | -9.2 | dbfda-world_1048 | | | Rifaximin | | -9.7 | | | DrugBank_approved_3 | | | Desmopressin |
| -9.5 | DrugBank_approved_2079 | Entrectinib | | -9.2 | dbfda-world_549 | | | Eplerenone | | -9.7 | | | dbfda-world_496 | | | Nystatin |
| -9.4 | DrugBank_approved_555 | Ergotamine | | -9.2 | dbfda-world_545 | | | Ergotamine | | -9.7 | | | dbfda-world_545 | | | Ergotamine |
| -9.4 | dbfda-world_456 | Ivermectin | | -9.2 | TheBindingDB_559 | | | Epinephrine | | -9.7 | | | dbfda-world_494 | | | Nystatin |
| -9.4 | dbfda-world_531 | Amphotericin B | | -9.2 | dbfda-world_1519 | | | Trypan blue | | -9.6 | | | dbfda-world_1534 | | | Paritaprevir |
| -9.4 | DrugBank_approved_540 | Amphotericin B | | -9.2 | dbfda-world_494 | | | Nystatin | | -9.6 | | | dbfda-world_456 | | | Ivermectin |
| -9.3 | TheBindingDB_821 | Camptosar | | -9.1 | dbfda-world_1534 | | | Paritaprevir | | -9.5 | | | dbfda-world_1629 | | | Naldemedine |
| -9.3 | dbfda-world_1628 | Velpatasvir | | -9.1 | dbfda-world_495 | | | Nystatin | | -9.5 | | | DrugBank_approved_1962 | | | Bisoctrizole |
| -9.2 | dbfda-world_545 | Ergotamine | | -9.1 | DrugBank_approved_3 | | | Desmopressin | | -9.5 | | | TheBindingDB_559 | | | B.H And Epinephrine |
| -9.2 | dbfda-world_1028 | Rifapentine | | -9.1 | dbfda-world_531 | | | Amphotericin B | | -9.4 | | | DrugBank_approved_90 | | | Adapalene |
|  |  |  | | -9.1 | ChemBridge_26 | | | 7786009 | | -9.4 | | | TheBindingDB_938 | | | Concentraid |
| **TMPRSS2_1** | | | | **TMPRSS2_2** | | | | | | **TMPRSS2_3** | | | | | | |
| **Energy** | **Database** | **Name** | | **Energy** | **Database** | | | **Name** | | **Energy** | | | **Database** | | | **Name** |
| -11.2 | DrugBank_approved_2108 | Cantharidin | | -11.8 | DrugBank_approved_980 | | | Candicidin | | -11.3 | | | DrugBank_approved_2005 | | | Venetoclax |
| -10.7 | dbfda-world_256 | Digoxin | | -10.7 | DrugBank_approved_2108 | | | Cantharidin | | -10.9 | | | DrugBank_approved_1137 | | | Quinupristin |
| -10.1 | DrugBank_approved_1137 | Quinupristin | | -10.2 | DrugBank_approved_811 | | | Dactinomycin | | -10.7 | | | DrugBank_approved_980 | | | Candicidin |
| -9.7 | DrugBank_approved_1401 | Icatibant | | -10.2 | DrugBank_approved_2005 | | | Venetoclax | | -10.4 | | | dbfda-world_176 | | | Dihydroergotamine |
| -9.6 | DrugBank_approved_506 | Nystatin | | -10.1 | dbfda-world_256 | | | Digoxin | | -10.3 | | | DrugBank_approved_1401 | | | Icatibant |
| -9.6 | DrugBank_approved_980 | Candicidin | | -10 | DrugBank_approved_486 | | | Bacitracin | | -10.3 | | | dbfda-world_531 | | | Amphotericin B |
| -9.5 | dbfda-world_531 | Amphotericin B | | -9.8 | dbfda-world_494 | | | Nystatin | | -10.3 | | | DrugBank_approved_2108 | | | Cantharidin |
| -9.4 | DrugBank_approved_555 | Ergotamine | | -9.7 | DrugBank_approved_1137 | | | Quinupristin | | -10.3 | | | dbfda-world_494 | | | Nystatin |
| -9.4 | dbfda-world_496 | Nystatin | | -9.6 | DrugBank_approved_1339 | | | Nilotinib | | -10.1 | | | DrugBank_approved_506 | | | Nystatin |
| -9.4 | dbfda-world_1263 | Eltrombopag | | -9.4 | DrugBank_approved_506 | | | Nystatin | | -10.1 | | | DrugBank_approved_377 | | | Acetyldigitoxin |
| -9.4 | dbfda-world_1519 | Trypan blue | | -9.4 | dbfda-world_531 | | | Amphotericin B | | -10 | | | DrugBank_approved_2441 | | | Ubrogepant |
| -9.3 | DrugBank_approved_2441 | Ubrogepant | | -9.4 | DrugBank_approved_1639 | | | Ledipasvir | | -10 | | | dbfda-world_1299 | | | Midostaurin |
| -9.3 | DrugBank_approved_955 | Dutasteride | | -9.3 | TheBindingDB_955 | | | Tasigna | | -10 | | | dbfda-world_256 | | | Digoxin |
| -9.2 | dbfda-world_493 | Nystatin | | -9.3 | DrugBank_approved_377 | | | Acetyldigitoxin | | -9.9 | | | dbfda-world_1534 | | | Paritaprevir |
| -9.2 | DrugBank_approved_1814 | Tannic acid | | -9.3 | TheBindingDB_910 | | | Avagard | | -9.9 | | | DrugBank_approved_3 | | | Desmopressin |
| -9.2 | DrugBank_approved_2179 | Plecanatide | | -9.3 | TheBindingDB_1061 | | | Vaprisol | | -9.9 | | | dbfda-world_1519 | | | Trypan blue |
| -9.1 | DrugBank_approved_2429 | Dexamethasone M | | -9.2 | dbfda-world_735 | | | Sirolimus | | -9.8 | | | DrugBank_approved_2429 | | | Dexamethasone M |
| -9.1 | dbfda-world_494 | Nystatin | | -9.2 | dbfda-world_496 | | | Nystatin | | -9.8 | | | dbfda-world_496 | | | Nystatin |
| -8.8 | DrugBank_approved_1057 | Saquinavir | | -9.1 | dbfda-world_1534 | | | Paritaprevir | | -9.8 | | | DrugBank_approved_1814 | | | Tannic acid |
|  |  |  | | -9.1 | TheBindingDB_1155 | | | Membraneblue | |  | | |  | | |  |
| **S1/S2_1** | | | **S1/S2_2** | | | | | | **S1/S2_3** | | | | | | | |
| **Energy** | **Database** | **Name** | | **Energy** | **Database** | | | **Name** | | **Energy** | | | **Database** | | | **Name** |
| -11.7 | DrugBank_approved_980 | Candicidin | | -11.1 | DrugBank_approved_980 | | | Candicidin | | -9.2 | | | dbfda-world_495 | | | Nystatin |
| -10.7 | dbfda-world_494 | Nystatin | | -10.1 | dbfda-world_1519 | | | Trypan blue | | -9 | | | DrugBank_approved_980 | | | Candicidin |
| -10.5 | dbfda-world_176 | Dihydroergotamine | | -9.5 | dbfda-world_256 | | | Digoxin | | -9 | | | dbfda-world_494 | | | Nystatin |
| -10.4 | DrugBank_approved_3 | Desmopressin | | -9.3 | dbfda-world_1048 | | | Rifaximin | | -8.9 | | | dbfda-world_256 | | | Digoxin |
| -10.3 | TheBindingDB_1156 | Evans Blue | | -9.2 | DrugBank_approved_2441 | | | Ubrogepant | | -8.7 | | | DrugBank_approved_555 | | | Ergotamine |
| -10.2 | DrugBank_approved_2441 | Ubrogepant | | -9.2 | dbfda-world_494 | | | Nystatin | | -8.5 | | | TheBindingDB_1061 | | | Vaprisol |
| -10.1 | DrugBank_approved_555 | Ergotamine | | -9.1 | DrugBank_approved_8 | | | Abarelix | | -8.5 | | | dbfda-world_1519 | | | Trypan blue |
| -10.1 | dbfda-world_256 | Digoxin | | -9 | dbfda-world_1627 | | | Lifitegrast | | -8.4 | | | DrugBank_approved_2079 | | | Entrectinib |
| -10 | DrugBank_approved_1639 | Ledipasvir | | -9 | TheBindingDB_625 | | | Cafergot | | -8.4 | | | DrugBank_approved_1814 | | | Tannic acid |
| -9.9 | DrugBank_approved_1814 | Tannic acid | | -9 | DrugBank_approved_2108 | | | Cantharidin | | -8.4 | | | DrugBank_approved_2005 | | | Venetoclax |
| -9.9 | TheBindingDB_1061 | Vaprisol | | -8.8 | dbfda-world_690 | | | Natamycin | | -8.3 | | | DrugBank_approved_506 | | | Nystatin |
| -9.9 | DrugBank_approved_2005 | Venetoclax | | -8.7 | dbfda-world_736 | | | Sirolimus | | -8.3 | | | DrugBank_approved_2056 | | | Capmatinib |
| -9.8 | DrugBank_approved_2082 | Avatrombopag | | -8.7 | dbfda-world_1117 | | | Ciclesonide | | -8.3 | | | dbfda-world_493 | | | Nystatin |
| -9.8 | DrugBank_approved_1736 | Trypan blue | | -8.7 | DrugBank_approved_618 | | | Irinotecan | | -8.3 | | | dbfda-world_496 | | | Nystatin |
| -9.8 | dbfda-world_531 | Amphotericin B | | -8.6 | DrugBank_approved_1137 | | | Quinupristin | | -8.3 | | | dbfda-world_531 | | | Amphotericin B |
| -9.8 | DrugBank_approved_479 | Imatinib | | -8.6 | dbfda-world_176 | | | Dihydroergotamine | | -8.3 | | | TheBindingDB_770 | | | Eptifibatide |
| -9.7 | dbfda-world_1283 | Simeprevir | | -8.6 | dbfda-world_496 | | | Nystatin | | -8.3 | | | DrugBank_approved_486 | | | Bacitracin |
| -9.7 | DrugBank_approved_506 | Nystatin | | -8.6 | DrugBank_approved_540 | | | Amphotericin B | | -8.2 | | | TheBindingDB_230 | | | Votrient |
| -9.7 | dbfda-world_1534 | Paritaprevir | | -8.6 | DrugBank_approved_302 | | | Cefpiramide | | -8.2 | | | dbfda-world_730 | | | Conivaptan |
| -9.7 | dbfda-world_495 | Nystatin | | -8.6 | DrugBank_approved_1639 | | | Ledipasvir | |  | | |  | | |  |
| **S1/S2'_1^+^** | | | **S1/S2'_2^+^** | | | | | | **S1/S2'_3^+^** | | | | | | | |
| **Energy** | **Database** | **Name** | | **Energy** | **Database** | | | **Name** | | **Energy** | | | **Database** | | | **Name** |
| -8.5 | DrugBank_approved_2108 | Cantharidin | | -9 | dbfda-world_256 | | | Digoxin | | -8.9 | | | dbfda-world_494 | | | Nystatin |
| -8.5 | dbfda-world_494 | Nystatin | | -8.4 | dbfda-world_176 | | | Dihydroergotamine | | -8.7 | | | dbfda-world_256 | | | Digoxin |
| -8.4 | dbfda-world_495 | Nystatin | | -8.4 | dbfda-world_496 | | | Nystatin | | -8.6 | | | DrugBank_approved_540 | | | Amphotericin B |
| -8.4 | DrugBank_approved_980 | Candicidin | | -8.2 | DrugBank_approved_980 | | | Candicidin | | -8.6 | | | DrugBank_approved_2108 | | | Cantharidin |
| -8.3 | dbfda-world_689 | Natamycin | | -8 | dbfda-world_690 | | | Natamycin | | -8.5 | | | DrugBank_approved_980 | | | Candicidin |
| -8.3 | dbfda-world_256 | Digoxin | | -7.9 | dbfda-world_1407 | | | Eribulin | | -8.2 | | | DrugBank_approved_3 | | | Desmopressin |
| -8.2 | dbfda-world_690 | Natamycin | | -7.9 | dbfda-world_545 | | | Ergotamine | | -8.2 | | | dbfda-world_736 | | | Sirolimus |
| -8.1 | dbfda-world_493 | Nystatin | | -7.9 | dbfda-world_1519 | | | Trypan blue | | -8.2 | | | DrugBank_approved_2005 | | | Venetoclax |
| -8.1 | dbfda-world_1519 | Trypan blue | | -7.8 | dbfda-world_1028 | | | Rifapentine | | -8.1 | | | dbfda-world_1048 | | | Rifaximin |
| -7.8 | DrugBank_approved_506 | Nystatin | | -7.7 | DrugBank_approved_2056 | | | Capmatinib | | -8 | | | DrugBank_approved_1137 | | | Quinupristin |
| -7.8 | dbfda-world_736 | Sirolimus | | -7.7 | dbfda-world_494 | | | Nystatin | | -8 | | | dbfda-world_495 | | | Nystatin |
| -7.8 | dbfda-world_531 | Amphotericin B | | -7.6 | DrugBank_approved_2079 | | | Entrectinib | | -7.9 | | | dbfda-world_1282 | | | Simeprevir |
| -7.7 | DrugBank_approved_1137 | Quinupristin | | -7.6 | DrugBank_approved_555 | | | Ergotamine | | -7.9 | | | dbfda-world_496 | | | Nystatin |
| -7.7 | dbfda-world_496 | Nystatin | | -7.6 | dbfda-world_1627 | | | Lifitegrast | | -7.9 | | | TheBindingDB_660 | | | Ivacaftor |
| -7.6 | dbfda-world_1283 | Simeprevir | | -7.6 | dbfda-world_531 | | | Amphotericin B | | -7.8 | | | DrugBank_approved_506 | | | Nystatin |
| -7.6 | DrugBank_approved_2005 | Venetoclax | | -7.6 | TheBindingDB_770 | | | Eptifibatide | | -7.8 | | | dbfda-world_689 | | | Natamycin |
| -7.6 | dbfda-world_545 | Ergotamine | | -7.6 | DrugBank_approved_2005 | | | Venetoclax | | -7.8 | | | dbfda-world_531 | | | Amphotericin B |
| -7.5 | dbfda-world_1573 | Ecamsule | | -7.5 | TheBindingDB_1035 | | | D.H.E. 45 | | -7.8 | | | dbfda-world_545 | | | Ergotamine |
|  |  |  | | -7.4 | dbfda-world_495 | | | Nystatin | | -7.7 | | | dbfda-world_1629 | | | Naldemedine |
|  |  |  | | -7.4 | DrugBank_approved_2441 | | | Ubrogepant | | -7.7 | | | DrugBank_approved_555 | | | Ergotamine |
| **S2'** | | | | **S2'_D936Y** | | | | | |  | | | | | | |
| **Energy** | **Database** | **Name** | | **Energy** | **Database** | | | **Name** | |  |  | | | |  | |
| -12.8 | DrugBank_approved_980 | Candicidin | | -10.4 | | DrugBank_approved_1137 | Quinupristin | | |  | |  | |  | | |
| -12.6 | dbfda-world_256 | Digoxin | | -10.3 | | dbfda-world_256 | Digoxin | | |  | |  | |  | | |
| -11.6 | DrugBank_approved_2005 | Venetoclax | | -10.2 | | DrugBank_approved_955 | Dutasteride | | |  | |  | |  | | |
| -11.4 | DrugBank_approved_2108 | Cantharidin | | -10.1 | | TheBindingDB_722 | Viracept | | |  | |  | |  | | |
| -11.2 | DrugBank_approved_1162 | Antrafenine | | -10.1 | | dbfda-world_953 | Atovaquone | | |  | |  | |  | | |
| -11 | TheBindingDB_780 | Juxtapid | | -9.9 | | dbfda-world_1477 | Olaparib | | |  | |  | |  | | |
| -10.9 | dbfda-world_1629 | Naldemedine | | -9.9 | | dbfda-world_1070 | Docetaxel | | |  | |  | |  | | |
| -10.9 | TheBindingDB_1285 | IC-Green | | -9.8 | | TheBindingDB_952 | Glyxambi | | |  | |  | |  | | |
| -10.8 | DrugBank_approved_377 | Acetyldigitoxin | | -9.8 | | dbfda-world_1282 | Simeprevir | | |  | |  | |  | | |
| -10.8 | DrugBank_approved_1565 | Lomitapide | | -9.8 | | TheBindingDB_689 | Natamycin | | |  | |  | |  | | |
| -10.7 | DrugBank_approved_1137 | Quinupristin | | -9.8 | | dbfda-world_1179 | Deoxycholic acid | | |  | |  | |  | | |
| -10.7 | TheBindingDB_408 | Vaprisol | | -9.8 | | TheBindingDB_1061 | Vaprisol | | |  | |  | |  | | |
| -10.6 | DrugBank_approved_1401 | Icatibant | | -9.8 | | DrugBank_approved_720 | Conivaptan | | |  | |  | |  | | |
| -10.6 | dbfda-world_1519 | Trypan blue | | -9.8 | | - | Drospirenone | | |  | |  | |  | | |
| -10.5 | DrugBank_approved_8 | Abarelix | | -9.7 | | DrugBank_approved_868 | Irbesartan | | |  | |  | |  | | |
| -10.3 | TheBindingDB_821 | Camptosar | | -9.7 | | DrugBank_approved_3 | Desmopressin | | |  | |  | |  | | |
| -10.3 | dbfda-world_656 | Candesartan cilexetil | | -9.7 | | DrugBank_approved_2058 | Bictegravir | | |  | |  | |  | | |
| -10.3 | dbfda-world_1081 | Posaconazole | | -9.7 | | dbfda-world_1058 | Saquinavir | | |  | |  | |  | | |
|  |  |  | | -9.7 | | TheBindingDB_688 | Irbesartan | | |  | |  | |  | | |

*Unit of energy is kcal/mol. AutoDock Vina (version 1.1.2, Linux) was used for the docking.

+ S1/S2’ corresponds to the cleavage site targeted by catL (T696-M697) on S1/S2 boundary.

**Table S2.** Top ~20 small molecule hits targeting the complexes obtained by *in silico* screening *

| **trypsin_1-S1/S2_1** | | | | | | | | | | | | | | | | | | | | | **trypsin_1-S1/S2_2** | | | | | | | | | | | | | | | | | | | | | | | **trypsin_2-S1/S2_1** | | | | | | | | | | | | | | | | | | | | | | | | | | | | | | | | | | | | |
| --- | --- | --- | --- | --- | --- | --- | --- | --- | --- | --- | --- | --- | --- | --- | --- | --- | --- | --- | --- | --- | --- | --- | --- | --- | --- | --- | --- | --- | --- | --- | --- | --- | --- | --- | --- | --- | --- | --- | --- | --- | --- | --- | --- | --- | --- | --- | --- | --- | --- | --- | --- | --- | --- | --- | --- | --- | --- | --- | --- | --- | --- | --- | --- | --- | --- | --- | --- | --- | --- | --- | --- | --- | --- | --- | --- | --- | --- | --- | --- | --- |
| **Energy** | | | | | | | **Database** | | | | | | | **Name** | | | | | | | **Energy** | | | | | | | | | | | **Database** | | | | | | | | **Name** | | | | **Energy** | | | | | | | | | | | | **Database** | | | | | | | | | | | | | | | | | | | | | | | | **Name** |
| -11.7 | | | | | | | dbfda-world_1263 | | | | | | | Eltrombopag | | | | | | | -11.5 | | | | | | | | | | | DrugBank_approved_555 | | | | | | | | Ergotamine | | | | -12.2 | | | | | | | | | | | | dbfda-world_1629 | | | | | | | | | | | | | | | | | | | | | | | | Naldemedine |
| -11.5 | | | | | | | DrugBank_approved_1773 | | | | | | | Lumacaftor | | | | | | | -11.4 | | | | | | | | | | | dbfda-world_1654 | | | | | | | | Glecaprevir | | | | -11.4 | | | | | | | | | | | | DrugBank_approved_1401 | | | | | | | | | | | | | | | | | | | | | | | | Icatibant |
| -11.4 | | | | | | | TheBindingDB_253 | | | | | | | Lapatinib | | | | | | | -11.4 | | | | | | | | | | | dbfda-world_1629 | | | | | | | | Naldemedine | | | | -11.2 | | | | | | | | | | | | DrugBank_approved_2005 | | | | | | | | | | | | | | | | | | | | | | | | Venetoclax |
| -11.3 | | | | | | | ChemBridge_26 | | | | | | | 7786009 | | | | | | | -11.4 | | | | | | | | | | | dbfda-world_256 | | | | | | | | Digoxin | | | | -11.1 | | | | | | | | | | | | DrugBank_approved_3 | | | | | | | | | | | | | | | | | | | | | | | | Desmopressin |
| -11.3 | | | | | | | dbfda-world_1572 | | | | | | | Ecamsule | | | | | | | -11.1 | | | | | | | | | | | dbfda-world_690 | | | | | | | | Natamycin | | | | -11.1 | | | | | | | | | | | | DrugBank_approved_980 | | | | | | | | | | | | | | | | | | | | | | | | Candicidin |
| -11.3 | | | | | | | DrugBank_approved_2155 | | | | | | | Pexidartinib | | | | | | | -11 | | | | | | | | | | | dbfda-world_1075 | | | | | | | | Retapamulin | | | | -11 | | | | | | | | | | | | dbfda-world_1519 | | | | | | | | | | | | | | | | | | | | | | | | Trypan blue |
| -11.2 | | | | | | | TheBindingDB_955 | | | | | | | Tasigna | | | | | | | -11 | | | | | | | | | | | DrugBank_approved_2056 | | | | | | | | Capmatinib | | | | -10.9 | | | | | | | | | | | | dbfda-world_496 | | | | | | | | | | | | | | | | | | | | | | | | Nystatin |
| -11 | | | | | | | TheBindingDB_565 | | | | | | | Nupercaine | | | | | | | -11 | | | | | | | | | | | dbfda-world_1070 | | | | | | | | Docetaxel | | | | -10.8 | | | | | | | | | | | | DrugBank_approved_2066 | | | | | | | | | | | | | | | | | | | | | | | | Revefenacin |
| -10.9 | | | | | | | DrugBank_approved_1162 | | | | | | | Antrafenine | | | | | | | -11 | | | | | | | | | | | dbfda-world_1028 | | | | | | | | Rifapentine | | | | -10.8 | | | | | | | | | | | | dbfda-world_256 | | | | | | | | | | | | | | | | | | | | | | | | Digoxin |
| -10.9 | | | | | | | DrugBank_approved_2441 | | | | | | | Ubrogepant | | | | | | | -10.9 | | | | | | | | | | | dbfda-world_176 | | | | | | | | Dihydroergotamine | | | | -10.7 | | | | | | | | | | | | DrugBank_approved_259 | | | | | | | | | | | | | | | | | | | | | | | | Valrubicin |
| -10.9 | | | | | | | DrugBank_approved_2056 | | | | | | | Capmatinib | | | | | | | -10.9 | | | | | | | | | | | dbfda-world_689 | | | | | | | | Natamycin | | | | -10.7 | | | | | | | | | | | | DrugBank_approved_1263 | | | | | | | | | | | | | | | | | | | | | | | | FAD |
| -10.9 | | | | | | | DrugBank_approved_1339 | | | | | | | Nilotinib | | | | | | | -10.9 | | | | | | | | | | | TheBindingDB_559 | | | | | | | | Epinephrine | | | | -10.6 | | | | | | | | | | | | TheBindingDB_955 | | | | | | | | | | | | | | | | | | | | | | | | Tasigna |
| -10.9 | | | | | | | TheBindingDB_529 | | | | | | | Thyroid Hormone | | | | | | | -10.9 | | | | | | | | | | | DrugBank_approved_980 | | | | | | | | Candicidin | | | | -10.5 | | | | | | | | | | | | dbfda-world_176 | | | | | | | | | | | | | | | | | | | | | | | | Dihydroergotamine |
| -10.9 | | | | | | | DrugBank_approved_1442 | | | | | | | Pazopanib | | | | | | | -10.8 | | | | | | | | | | | dbfda-world_545 | | | | | | | | Ergotamine | | | | -10.5 | | | | | | | | | | | | DrugBank_approved_1339 | | | | | | | | | | | | | | | | | | | | | | | | Nilotinib |
| -10.8 | | | | | | | DrugBank_approved_2005 | | | | | | | Venetoclax | | | | | | | -10.7 | | | | | | | | | | | dbfda-world_1534 | | | | | | | | Paritaprevir | | | | -10.5 | | | | | | | | | | | | dbfda-world_730 | | | | | | | | | | | | | | | | | | | | | | | | Conivaptan |
| -10.8 | | | | | | | TheBindingDB_572 | | | | | | | Droperidol | | | | | | | -10.7 | | | | | | | | | | | TheBindingDB_625 | | | | | | | | Cafergot | | | | -10.5 | | | | | | | | | | | | DrugBank_approved_1814 | | | | | | | | | | | | | | | | | | | | | | | | Tannic acid |
| -10.7 | | | | | | | DrugBank_approved_1652 | | | | | | | Ibrutinib | | | | | | | -10.7 | | | | | | | | | | | dbfda-world_1519 | | | | | | | | Trypan blue | | | | -10.5 | | | | | | | | | | | | dbfda-world_1428 | | | | | | | | | | | | | | | | | | | | | | | | Ponatinib |
| -10.7 | | | | | | | DrugBank_approved_1814 | | | | | | | Tannic Acid | | | | | | | -10.6 | | | | | | | | | | | dbfda-world_1621 | | | | | | | | Venetoclax | | | | -10.5 | | | | | | | | | | | | DrugBank_approved_1632 | | | | | | | | | | | | | | | | | | | | | | | | Diosmin |
| -10.7 | | | | | | | DrugBank_approved_2058 | | | | | | | Bictegravir | | | | | | | -10.6 | | | | | | | | | | | dbfda-world_456 | | | | | | | | Ivermectin | | | | -10.5 | | | | | | | | | | | | dbfda-world_545 | | | | | | | | | | | | | | | | | | | | | | | | Ergotamine |
| -10.7 | | | | | | | ChemBridge_29 | | | | | | | 7928214 | | | | | | | -10.6 | | | | | | | | | | | dbfda-world_1074 | | | | | | | | Retapamulin | | | | -10.3 | | | | | | | | | | | | dbfda-world_1217 | | | | | | | | | | | | | | | | | | | | | | | | Nilotinib |
| -10.7 | | | | | | | DrugBank_approved_1543 | | | | | | | Triptorelin | | | | | | | -10.5 | | | | | | | | | | | DrugBank_approved_377 | | | | | | | | Acetyldigitoxin | | | | -10.3 | | | | | | | | | | | | TheBindingDB_232 | | | | | | | | | | | | | | | | | | | | | | | | Gleevec |
| -10.7 | | | | | | | TheBindingDB_780 | | | | | | | Juxtapid | | | | | | | -10.5 | | | | | | | | | | | - | | | | | | | | Camostat | | | | -10.3 | | | | | | | | | | | | DrugBank_approved_2108 | | | | | | | | | | | | | | | | | | | | | | | | Cantharidin |
| -9.5 | | | | | | | - | | | | | | | Camostat | | | | | | |  | | | | | | | | | | |  | | | | | | | |  | | | |  | | | | | | | | | | | | | |  | | | | | | | | | | | | | | | | | | | | | | |
| **trypsin_1-S1/S2_2** | | | | | | | | | | | | | | | | | | | | | **trypsin_2-S1/S2_2** | | | | | | | | | | | | | | | | | | | | | | | **trypsin_3-S1/S2_2** | | | | | | | | | | | | | | | | | | | | | | | | | | | | | | | | | | | | |
| **Energy** | | | **Database** | | | | | | | **Name** | | | | | | | **Energy** | | | | | | | | | | **Database** | | | | | | | | **Name** | | | | | | | | | **Energy** | | | | | | | | | | | **Database** | | | | | | | | | | | | | | | | | | | | | | | **Name** | | |
| -12.2 | | | DrugBank_approved_980 | | | | | | | Candicidin | | | | | | | -11.3 | | | | | | | | | | DrugBank_approved_1137 | | | | | | | | Quinupristin | | | | | | | | | -11 | | | | | | | | | | | DrugBank_approved_980 | | | | | | | | | | | | | | | | | | | | | | | Candicidin | | |
| -11.6 | | | dbfda-world_494 | | | | | | | Nystatin | | | | | | | -11.3 | | | | | | | | | | DrugBank_approved_980 | | | | | | | | Candicidin | | | | | | | | | -10.7 | | | | | | | | | | | dbfda-world_256 | | | | | | | | | | | | | | | | | | | | | | | Digoxin | | |
| -10.9 | | | dbfda-world_256 | | | | | | | Digoxin | | | | | | | -11.2 | | | | | | | | | | dbfda-world_256 | | | | | | | | Digoxin | | | | | | | | | -10.5 | | | | | | | | | | | dbfda-world_496 | | | | | | | | | | | | | | | | | | | | | | | Nystatin | | |
| -10.4 | | | dbfda-world_493 | | | | | | | Nystatin | | | | | | | -11 | | | | | | | | | | dbfda-world_176 | | | | | | | | Dihydroergotamine | | | | | | | | | -10.4 | | | | | | | | | | | dbfda-world_495 | | | | | | | | | | | | | | | | | | | | | | | Nystatin | | |
| -10.4 | | | DrugBank_approved_2108 | | | | | | | Cantharidin | | | | | | | -11 | | | | | | | | | | DrugBank_approved_1814 | | | | | | | | Tannic acid | | | | | | | | | -10.4 | | | | | | | | | | | DrugBank_approved_2108 | | | | | | | | | | | | | | | | | | | | | | | Cantharidin | | |
| -10.3 | | | DrugBank_approved_506 | | | | | | | Nystatin | | | | | | | -10.9 | | | | | | | | | | DrugBank_approved_720 | | | | | | | | Conivaptan | | | | | | | | | -10.3 | | | | | | | | | | | dbfda-world_1519 | | | | | | | | | | | | | | | | | | | | | | | Trypan blue | | |
| -9.9 | | | DrugBank_approved_3 | | | | | | | Desmopressin | | | | | | | -10.8 | | | | | | | | | | dbfda-world_1621 | | | | | | | | Venetoclax | | | | | | | | | -10.3 | | | | | | | | | | | dbfda-world_494 | | | | | | | | | | | | | | | | | | | | | | | Nystatin | | |
| -9.9 | | | dbfda-world_531 | | | | | | | Amphotericin B | | | | | | | -10.8 | | | | | | | | | | TheBindingDB_955 | | | | | | | | Tasigna | | | | | | | | | -10.1 | | | | | | | | | | | DrugBank_approved_506 | | | | | | | | | | | | | | | | | | | | | | | Nystatin | | |
| -9.8 | | | dbfda-world_176 | | | | | | | Dihydroergotamine | | | | | | | -10.8 | | | | | | | | | | dbfda-world_1534 | | | | | | | | Paritaprevir | | | | | | | | | -10 | | | | | | | | | | | dbfda-world_1534 | | | | | | | | | | | | | | | | | | | | | | | Paritaprevir | | |
| -9.8 | | | dbfda-world_496 | | | | | | | Nystatin | | | | | | | -10.8 | | | | | | | | | | DrugBank_approved_1339 | | | | | | | | Nilotinib | | | | | | | | | -9.7 | | | | | | | | | | | dbfda-world_493 | | | | | | | | | | | | | | | | | | | | | | | Nystatin | | |
| -9.8 | | | DrugBank_approved_1401 | | | | | | | Icatibant | | | | | | | -10.8 | | | | | | | | | | ChemBridge_26 | | | | | | | | 7786009 | | | | | | | | | -9.7 | | | | | | | | | | | DrugBank_approved_486 | | | | | | | | | | | | | | | | | | | | | | | Bacitracin | | |
| -9.8 | | | dbfda-world_1519 | | | | | | | Trypan blue | | | | | | | -10.7 | | | | | | | | | | DrugBank_approved_2005 | | | | | | | | Venetoclax | | | | | | | | | -9.6 | | | | | | | | | | | dbfda-world_1299 | | | | | | | | | | | | | | | | | | | | | | | Midostaurin | | |
| -9.6 | | | DrugBank_approved_1137 | | | | | | | Quinupristin | | | | | | | -10.6 | | | | | | | | | | dbfda-world_1217 | | | | | | | | Nilotinib | | | | | | | | | -9.6 | | | | | | | | | | | dbfda-world_531 | | | | | | | | | | | | | | | | | | | | | | | Amphotericin B | | |
| -9.6 | | | dbfda-world_1534 | | | | | | | Paritaprevir | | | | | | | -10.6 | | | | | | | | | | TheBindingDB_71 | | | | | | | | Ritonavir | | | | | | | | | -9.5 | | | | | | | | | | | dbfda-world_176 | | | | | | | | | | | | | | | | | | | | | | | Dihydroergotamine | | |
| -9.6 | | | dbfda-world_495 | | | | | | | Nystatin | | | | | | | -10.5 | | | | | | | | | | dbfda-world_75 | | | | | | | | Adapalene | | | | | | | | | -9.4 | | | | | | | | | | | dbfda-world_1654 | | | | | | | | | | | | | | | | | | | | | | | Glecaprevir | | |
| -9.6 | | | DrugBank_approved_486 | | | | | | | Bacitracin | | | | | | | -10.4 | | | | | | | | | | TheBindingDB_568 | | | | | | | | Vontrol | | | | | | | | | -9.4 | | | | | | | | | | | dbfda-world_1477 | | | | | | | | | | | | | | | | | | | | | | | Olaparib | | |
| -9.5 | | | DrugBank_approved_5 | | | | | | | Daptomycin | | | | | | | -10.4 | | | | | | | | | | ChemBridge_10 | | | | | | | | 7793889 | | | | | | | | | -9.4 | | | | | | | | | | | TheBindingDB_299 | | | | | | | | | | | | | | | | | | | | | | | Dactinomycin | | |
| -9.5 | | | dbfda-world_1028 | | | | | | | Rifapentine | | | | | | | -10.3 | | | | | | | | | | DrugBank_approved_2079 | | | | | | | | Entrectinib | | | | | | | | | -9.4 | | | | | | | | | | | TheBindingDB_1320 | | | | | | | | | | | | | | | | | | | | | | | Tannic Acid | | |
| -9.4 | | | DrugBank_approved_1028 | | | | | | | Rifapentine | | | | | | | -10.3 | | | | | | | | | | TheBindingDB_1229 | | | | | | | | Vemurafenib | | | | | | | | | -9.4 | | | | | | | | | | | TheBindingDB_278 | | | | | | | | | | | | | | | | | | | | | | | Rapamune | | |
| -9.3 | | | DrugBank_approved_540 | | | | | | | Amphotericin B | | | | | | | -10.3 | | | | | | | | | | DrugBank_approved_506 | | | | | | | | Nystatin | | | | | | | | | -9.3 | | | | | | | | | | | dbfda-world_1283 | | | | | | | | | | | | | | | | | | | | | | | Simeprevir | | |
| -9.3 | | | DrugBank_approved_2021 | | | | | | | Velpatasvir | | | | | | | -10.3 | | | | | | | | | | DrugBank_approved_555 | | | | | | | | Ergotamine | | | | | | | | | -9.3 | | | | | | | | | | | dbfda-world_1455 | | | | | | | | | | | | | | | | | | | | | | | Ledipasvir | | |
| -9.2 | | | DrugBank_approved_1814 | | | | | | | Tannic acid | | | | | | |  | | | | | | | | | |  | | | | | | | |  | | | | | | | | | -9.3 | | | | | | | | | | | TheBindingDB_625 | | | | | | | | | | | | | | | | | | | | | | | Cafergot | | |
| **trypsin_1-S2'** | | | | | | | | | | | | | | | | | | | | | **trypsin_2-S2'** | | | | | | | | | | | | | | | | | | | | | | | **trypsin_3-S2'** | | | | | | | | | | | | | | | | | | | | | | | | | | | | | | | | | | | | |
| **Energy** | | | | | **Database** | | | | | | | **Name** | | | | | | | | **Energy** | | | | | | | | | **Database** | | | | | | | | **Name** | | | | | | | **Energy** | | | | | | | | | **Database** | | | | | | | | | | | | | | | | | | | | | **Name** | | | | | | |
| -11.4 | | | | | DrugBank_approved_2108 | | | | | | | Cantharidin | | | | | | | | -12.8 | | | | | | | | | DrugBank_approved_2108 | | | | | | | | Cantharidin | | | | | | | -11.7 | | | | | | | | | DrugBank_approved_1137 | | | | | | | | | | | | | | | | | | | | | Quinupristin | | | | | | |
| -11.4 | | | | | dbfda-world_494 | | | | | | | Nystatin | | | | | | | | -11 | | | | | | | | | DrugBank_approved_506 | | | | | | | | Nystatin | | | | | | | -11.6 | | | | | | | | | DrugBank_approved_980 | | | | | | | | | | | | | | | | | | | | | Candicidin | | | | | | |
| -11.2 | | | | | TheBindingDB_760 | | | | | | | Prelay | | | | | | | | -10.6 | | | | | | | | | DrugBank_approved_980 | | | | | | | | Candicidin | | | | | | | -11 | | | | | | | | | dbfda-world_256 | | | | | | | | | | | | | | | | | | | | | Digoxin | | | | | | |
| -11.1 | | | | | DrugBank_approved_1162 | | | | | | | Antrafenine | | | | | | | | -10.2 | | | | | | | | | dbfda-world_256 | | | | | | | | Digoxin | | | | | | | -10.9 | | | | | | | | | dbfda-world_1572 | | | | | | | | | | | | | | | | | | | | | Ecamsule | | | | | | |
| -11 | | | | | TheBindingDB_955 | | | | | | | Tasigna | | | | | | | | -10.2 | | | | | | | | | dbfda-world_494 | | | | | | | | Nystatin | | | | | | | -10.5 | | | | | | | | | DrugBank_approved_506 | | | | | | | | | | | | | | | | | | | | | Nystatin | | | | | | |
| -11 | | | | | TheBindingDB_110 | | | | | | | Alimta | | | | | | | | -10.1 | | | | | | | | | DrugBank_approved_3 | | | | | | | | Desmopressin | | | | | | | -10.4 | | | | | | | | | TheBindingDB_1061 | | | | | | | | | | | | | | | | | | | | | Vapsirol | | | | | | |
| -10.9 | | | | | DrugBank_approved_2045 | | | | | | | Lasmiditan | | | | | | | | -10 | | | | | | | | | dbfda-world_1654 | | | | | | | | Glecaprevir | | | | | | | -10.2 | | | | | | | | | dbfda-world_1621 | | | | | | | | | | | | | | | | | | | | | Venetoclax | | | | | | |
| -10.9 | | | | | DrugBank_approved_2074 | | | | | | | Duvelisib | | | | | | | | -9.9 | | | | | | | | | DrugBank_approved_555 | | | | | | | | Ergotamine | | | | | | | -10.2 | | | | | | | | | DrugBank_approved_1773 | | | | | | | | | | | | | | | | | | | | | Lumacaftor | | | | | | |
| -10.8 | | | | | DrugBank_approved_946 | | | | | | | Atovaquone | | | | | | | | -9.8 | | | | | | | | | DrugBank_approved_1137 | | | | | | | | Quinupristin | | | | | | | -10.1 | | | | | | | | | DrugBank_approved_1814 | | | | | | | | | | | | | | | | | | | | | Tannic acid | | | | | | |
| -10.7 | | | | | ChemBridge_25 | | | | | | | 7931205 | | | | | | | | -9.8 | | | | | | | | | TheBindingDB_278 | | | | | | | | Rapamune | | | | | | | -10.1 | | | | | | | | | DrugBank_approved_1639 | | | | | | | | | | | | | | | | | | | | | Ledipasvir | | | | | | |
| -10.7 | | | | | dbfda-world_1654 | | | | | | | Glecaprevir | | | | | | | | -9.8 | | | | | | | | | TheBindingDB_873 | | | | | | | | Viibryd | | | | | | | -10.1 | | | | | | | | | DrugBank_approved_2005 | | | | | | | | | | | | | | | | | | | | | Venetoclax | | | | | | |
| -10.7 | | | | | DrugBank_approved_1814 | | | | | | | Tannic acid | | | | | | | | -9.7 | | | | | | | | | dbfda-world_495 | | | | | | | | Nystatin | | | | | | | -10 | | | | | | | | | TheBindingDB_938 | | | | | | | | | | | | | | | | | | | | | Concentraid | | | | | | |
| -10.7 | | | | | dbfda-world_256 | | | | | | | Digoxin | | | | | | | | -9.7 | | | | | | | | | DrugBank_approved_2002 | | | | | | | | Elbasvir | | | | | | | -10 | | | | | | | | | DrugBank_approved_555 | | | | | | | | | | | | | | | | | | | | | Ergotamine | | | | | | |
| -10.7 | | | | | dbfda-world_1519 | | | | | | | Trypan blue | | | | | | | | -9.7 | | | | | | | | | dbfda-world_1192 | | | | | | | | Estrone sulfate | | | | | | | -10 | | | | | | | | | dbfda-world_493 | | | | | | | | | | | | | | | | | | | | | Nystatin | | | | | | |
| -10.7 | | | | | DrugBank_approved_101 | | | | | | | Glimepiride | | | | | | | | -9.7 | | | | | | | | | TheBindingDB_174 | | | | | | | | Raltegravir | | | | | | | -10 | | | | | | | | | DrugBank_approved_486 | | | | | | | | | | | | | | | | | | | | | Bacitracin | | | | | | |
| -10.6 | | | | | DrugBank_approved_2079 | | | | | | | Entrectinib | | | | | | | | -9.7 | | | | | | | | | dbfda-world_1519 | | | | | | | | Trypan blue | | | | | | | -9.9 | | | | | | | | | dbfda-world_496 | | | | | | | | | | | | | | | | | | | | | Nystatin | | | | | | |
| -10.6 | | | | | dbfda-world_496 | | | | | | | Nystatin | | | | | | | | -9.6 | | | | | | | | | DrugBank_approved_955 | | | | | | | | Dutasteride | | | | | | | -9.9 | | | | | | | | | TheBindingDB_947 | | | | | | | | | | | | | | | | | | | | | Aprepitant | | | | | | |
| -10.5 | | | | | dbfda-world_495 | | | | | | | Nystatin | | | | | | | | -9.6 | | | | | | | | | dbfda-world_1306 | | | | | | | | Axitinib | | | | | | | -9.9 | | | | | | | | | DrugBank_approved_720 | | | | | | | | | | | | | | | | | | | | | Conivaptan | | | | | | |
| -10.5 | | | | | ChemBridge_37 | | | | | | | 7927879 | | | | | | | | -9.6 | | | | | | | | | TheBindingDB_1061 | | | | | | | | Vapsirol | | | | | | | -9.9 | | | | | | | | | dbfda-world_494 | | | | | | | | | | | | | | | | | | | | | Nystatin | | | | | | |
| -10.4 | | | | | dbfda-world_176 | | | | | | | Dihydroergotamine | | | | | | | | -9.6 | | | | | | | | | dbfda-world_1133 | | | | | | | | Ursodeoxycholic A. | | | | | | | -9.8 | | | | | | | | | dbfda-world_1534 | | | | | | | | | | | | | | | | | | | | | Paritaprevir | | | | | | |
| **trypsin_1-S2'_D936Y** | | | | | | | | | | | | | | | | | | | | | **trypsin_2-S2'_D936Y** | | | | | | | | | | | | | | | | | | | | | | | **trypsin_3-S2'_D936Y** | | | | | | | | | | | | | | | | | | | | | | | | | | | | | | | | | | | | |
| **Energy** | | | | | **Database** | | | | | | | **Name** | | | | | | | | **Energy** | | | | | | | | | **Database** | | | | | | | | **Name** | | | | | | | **Energy** | | | | | | | | | **Database** | | | | | | | | | | | | | | | | | | | | | | | **Name** | | | | |
| -12.2 | | | | | DrugBank_approved_980 | | | | | | | Candicidin | | | | | | | | -12 | | | | | | | | | dbfda-world_1572 | | | | | | | | Ecamsule | | | | | | | -12.9 | | | | | | | | | DrugBank_approved_980 | | | | | | | | | | | | | | | | | | | | | | | Candicidin | | | | |
| -11.4 | | | | | DrugBank_approved_1814 | | | | | | | Tannic acid | | | | | | | | -11.7 | | | | | | | | | dbfda-world_494 | | | | | | | | Nystatin | | | | | | | -12.6 | | | | | | | | | dbfda-world_256 | | | | | | | | | | | | | | | | | | | | | | | Digoxin | | | | |
| -11.4 | | | | | DrugBank_approved_2108 | | | | | | | Cantharidin | | | | | | | | -11.3 | | | | | | | | | DrugBank_approved_980 | | | | | | | | Candicidin | | | | | | | -12.3 | | | | | | | | | DrugBank_approved_1137 | | | | | | | | | | | | | | | | | | | | | | | Quinupristin | | | | |
| -10.9 | | | | | ChemBridge_26 | | | | | | | 7786009 | | | | | | | | -11.1 | | | | | | | | | DrugBank_approved_506 | | | | | | | | Nystatin | | | | | | | -11.6 | | | | | | | | | DrugBank_approved_2005 | | | | | | | | | | | | | | | | | | | | | | | Venetoclax | | | | |
| -10.8 | | | | | TheBindingDB_568 | | | | | | | Vontrol | | | | | | | | -11.1 | | | | | | | | | dbfda-world_730 | | | | | | | | Conivaptan | | | | | | | -11.6 | | | | | | | | | dbfda-world_1519 | | | | | | | | | | | | | | | | | | | | | | Trypan blue | | | | | |
| -10.8 | | | | | dbfda-world_495 | | | | | | | Nystatin | | | | | | | | -11 | | | | | | | | | TheBindingDB_695 | | | | | | | | Adapalene | | | | | | | -11.5 | | | | | | | | | dbfda-world_176 | | | | | | | | | | | | | | | | | | | | | | Dihydroergotamine | | | | | |
| -10.8 | | | | | DrugBank_approved_1776 | | | | | | | Rolapitant | | | | | | | | -10.9 | | | | | | | | | dbfda-world_1085 | | | | | | | | Paliperidone | | | | | | | -11.5 | | | | | | | | | dbfda-world_493 | | | | | | | | | | | | | | | | | | | | | | Nystatin | | | | | |
| -10.8 | | | | | dbfda-world_256 | | | | | | | Digoxin | | | | | | | | -10.9 | | | | | | | | | DrugBank_approved_8 | | | | | | | | Abarelix | | | | | | | -11.5 | | | | | | | | | DrugBank_approved_8 | | | | | | | | | | | | | | | | | | | | | | Abarelix | | | | | |
| -10.6 | | | | | DrugBank_approved_532 | | | | | | | Aprepitant | | | | | | | | -10.8 | | | | | | | | | DrugBank_approved_125 | | | | | | | | Ziprasidone | | | | | | | -11.4 | | | | | | | | | DrugBank_approved_2108 | | | | | | | | | | | | | | | | | | | | | | Cantharidin | | | | | |
| -10.6 | | | | | DrugBank_approved_1137 | | | | | | | Quinupristin | | | | | | | | -10.8 | | | | | | | | | dbfda-world_496 | | | | | | | | Nystatin | | | | | | | -11.2 | | | | | | | | | DrugBank_approved_555 | | | | | | | | | | | | | | | | | | | | | | Ergotamine | | | | | |
| -10.5 | | | | | DrugBank_approved_3 | | | | | | | Desmopressin | | | | | | | | -10.8 | | | | | | | | | dbfda-world_256 | | | | | | | | Digoxin | | | | | | | -11.2 | | | | | | | | | dbfda-world_495 | | | | | | | | | | | | | | | | | | | | | | Nystatin | | | | | |
| -10.5 | | | | | DrugBank_approved_1361 | | | | | | | Indacaterol | | | | | | | | -10.7 | | | | | | | | | dbfda-world_1627 | | | | | | | | Lifitegrast | | | | | | | -11.2 | | | | | | | | | dbfda-world_1282 | | | | | | | | | | | | | | | | | | | | | | Simeprevir | | | | | |
| -10.5 | | | | | DrugBank_approved_2005 | | | | | | | Venetoclax | | | | | | | | -10.7 | | | | | | | | | dbfda-world_1084 | | | | | | | | Paliperidone | | | | | | | -11.2 | | | | | | | | | TheBindingDB_1320 | | | | | | | | | | | | | | | | | | | | | | Tannic Acid | | | | | |
| -10.4 | | | | | DrugBank_approved_506 | | | | | | | Nystatin | | | | | | | | -10.6 | | | | | | | | | dbfda-world_22 | | | | | | | | Ergocalciferol | | | | | | | -11.1 | | | | | | | | | dbfda-world_531 | | | | | | | | | | | | | | | | | | | | | | Amphotericin B | | | | | |
| -10.4 | | | | | DrugBank_approved_1409 | | | | | | | Eltrombopag | | | | | | | | -10.6 | | | | | | | | | DrugBank_approved_590 | | | | | | | | Risperidone | | | | | | | -11.0 | | | | | | | | | dbfda-world_1075 | | | | | | | | | | | | | | | | | | | | | | Retapamulin | | | | | |
| -10.4 | | | | | DrugBank_approved_2056 | | | | | | | Capmatinib | | | | | | | | -10.6 | | | | | | | | | TheBindingDB_989 | | | | | | | | Promacta | | | | | | | -10.9 | | | | | | | | | dbfda-world_1629 | | | | | | | | | | | | | | | | | | | | | | Velpatasvir | | | | | |
| -10.4 | | | | | dbfda-world_496 | | | | | | | Nystatin | | | | | | | | -10.6 | | | | | | | | | TheBindingDB_174 | | | | | | | | Raltegravir | | | | | | | -10.9 | | | | | | | | | dbfda-world_1534 | | | | | | | | | | | | | | | | | | | | | | Paritaprevir | | | | | |
| -10.4 | | | | | DrugBank_approved_1669 | | | | | | | Olaparib | | | | | | | | -10.5 | | | | | | | | | dbfda-world_495 | | | | | | | | Nystatin | | | | | | | -10.9 | | | | | | | | | DrugBank_approved_377 | | | | | | | | | | | | | | | | | | | | | | Acetyldigitoxin | | | | | |
| -10.3 | | | | | dbfda-world_1212 | | | | | | | Nebivolol | | | | | | | | -10.5 | | | | | | | | | dbfda-world_954 | | | | | | | | Atovaquone | | | | | | | -10.8 | | | | | | | | | dbfda-world_730 | | | | | | | | | | | | | | | | | | | | | | Conivaptan | | | | | |
| -10.3 | | | | | DrugBank_approved_2274 | | | | | | | Netarsudil | | | | | | | |  | | | | | | | | |  | | | | | | | |  | | | | | | | -10.8 | | | | | | | | | TheBindingDB_332 | | | | | | | | | | | | | | | | | | | | | | Tannic acid | | | | | |
| **TMPRSS2_1-S1/S2_1** | | | | | | | | | | | | | | | | | | | | | **TMPRSS2_2-S1/S2_1** | | | | | | | | | | | | | | | | | | | | | | |  | | | | | | | | | | | | | | | | |  | | |  | | | | | | | | | | | | | | | | |
| **Energy** | | | | | **Database** | | | | | | | **Name** | | | | | | | | **Energy** | | | | | | | | | **Database** | | | | | | | | **Name** | | | | | | |  | | | | | | | | | | | | | | | | | |  | | | | | | | | | | | | | | | | | | |
| -13.6 | | | | | dbfda-world_256 | | | | | | | Digoxin | | | | | | | | -11.7 | | | | | | | | | dbfda-world_496 | | | | | | | | Nystatin | | | | | | |  | | | | |  | | | | | | | | | | | | | | | | | | | | |  | | | | | | | | | | |
| -13.2 | | | | | dbfda-world_531 | | | | | | | Amphotericin B | | | | | | | | -11.6 | | | | | | | | | dbfda-world_256 | | | | | | | | Digoxin | | | | | | |  | | | | |  | | | | | | | | | | | | | | | | | | | | |  | | | | | | | | | | |
| -12.8 | | | | | DrugBank_approved_1137 | | | | | | | Quinupristin | | | | | | | | -11.1 | | | | | | | | | dbfda-world_1534 | | | | | | | | Paritaprevir | | | | | | |  | | | | |  | | | | | | | | | | | | | | | | | | | | |  | | | | | | | | | | |
| -12.6 | | | | | TheBindingDB_299 | | | | | | | Dactinomycin | | | | | | | | -11.1 | | | | | | | | | TheBindingDB_332 | | | | | | | | Tannic acid | | | | | | |  | | | | |  | | | | | | | | | | | | | | | | | | | | |  | | | | | | | | | | |
| -12.5 | | | | | dbfda-world_493 | | | | | | | Nystatin | | | | | | | | -11 | | | | | | | | | DrugBank_approved_526 | | | | | | | | Nafarelin | | | | | | |  | | | | |  | | | | | | | | | | | | | | | | | | | | |  | | | | | | | | | | |
| -12.5 | | | | | dbfda-world_496 | | | | | | | Nystatin | | | | | | | | -10.9 | | | | | | | | | DrugBank_approved_377 | | | | | | | | Acetyldigitoxin | | | | | | |  | | | | |  | | | | | | | | | | | | | | | | | | | | |  | | | | | | | | | | |
| -12.4 | | | | | DrugBank_approved_811 | | | | | | | Dactinomycin | | | | | | | | -10.9 | | | | | | | | | DrugBank_approved_2005 | | | | | | | | Venetoclax | | | | | | |  | | | | |  | | | | | | | | | | | | | | | | | | | | |  | | | | | | | | | | |
| -12.4 | | | | | dbfda-world_494 | | | | | | | Nystatin | | | | | | | | -10.8 | | | | | | | | | dbfda-world_809 | | | | | | | | Telmisartan | | | | | | |  | | | | |  | | | | | | | | | | | | | | | | | | | | |  | | | | | | | | | | |
| -12.1 | | | | | dbfda-world_495 | | | | | | | Nystatin | | | | | | | | -10.8 | | | | | | | | | DrugBank_approved_1543 | | | | | | | | Triptorelin | | | | | | |  | | | | |  | | | | | | | | | | | | | | | | | | | | |  | | | | | | | | | | |
| -12 | | | | | DrugBank_approved_506 | | | | | | | Nystatin | | | | | | | | -10.8 | | | | | | | | | DrugBank_approved_980 | | | | | | | | Candicidin | | | | | | |  | | | | |  | | | | | | | | | | | | | | | | | | | | |  | | | | | | | | | | |
| -12 | | | | | DrugBank_approved_1814 | | | | | | | Tannic acid | | | | | | | | -10.7 | | | | | | | | | TheBindingDB_938 | | | | | | | | Concentraid | | | | | | |  | | | | |  | | | | | | | | | | | | | | | | | | | | |  | | | | | | | | | | |
| -12 | | | | | DrugBank_approved_980 | | | | | | | Candicidin | | | | | | | | -10.7 | | | | | | | | | DrugBank_approved_555 | | | | | | | | Ergotamine | | | | | | |  | | | | |  | | | | | | | | | | | | | | | | | | | | |  | | | | | | | | | | |
| -11.8 | | | | | dbfda-world_1519 | | | | | | | Trypan blue | | | | | | | | -10.7 | | | | | | | | | dbfda-world_176 | | | | | | | | Dihydroergotamine | | | | | | |  | | | | |  | | | | | | | | | | | | | | | | | | | | |  | | | | | | | | | | |
| -11.6 | | | | | DrugBank_approved_2005 | | | | | | | Venetoclax | | | | | | | | -10.7 | | | | | | | | | dbfda-world_1281 | | | | | | | | Simeprevir | | | | | | |  | | | | |  | | | | | | | | | | | | | | | | | | | | |  | | | | | | | | | | |
| -11.6 | | | | | dbfda-world_545 | | | | | | | Ergotamine | | | | | | | | -10.7 | | | | | | | | | dbfda-world_494 | | | | | | | | Nystatin | | | | | | |  | | | | |  | | | | | | | | | | | | | | | | | | | | |  | | | | | | | | | | |
| -11.4 | | | | | dbfda-world_1534 | | | | | | | Paritaprevir | | | | | | | | -10.6 | | | | | | | | | dbfda-world_1299 | | | | | | | | Midostaurin | | | | | | |  | | | | |  | | | | | | | | | | | | | | | | | | | | |  | | | | | | | | | | |
| -11.4 | | | | | TheBindingDB_1156 | | | | | | | Evans Blue | | | | | | | | -10.6 | | | | | | | | | DrugBank_approved_1639 | | | | | | | | Ledipasvir | | | | | | |  | | | | |  | | | | | | | | | | | | | | | | | | | | |  | | | | | | | | | | |
| -11.4 | | | | | TheBindingDB_770 | | | | | | | Eptifibatide | | | | | | | | -10.6 | | | | | | | | | DrugBank_approved_2002 | | | | | | | | Elbasvir | | | | | | |  | | | | |  | | | | | | | | | | | | | | | | | | | | |  | | | | | | | | | | |
| -11.3 | | | | | dbfda-world_176 | | | | | | | Dihydroergotamine | | | | | | | |  | | | | | | | | |  | | | | | | | |  | | | | | | |  | | | | |  | | | | | | | | | | | | | | | | | | | | |  | | | | | | | | | | |
| -11.3 | | | | | DrugBank_approved_1401 | | | | | | | Icatibant | | | | | | | |  | | | | | | | | |  | | | | | | | |  | | | | | | |  | | | | | | |  | | | | | | | | | | | | | | | | | | | | | |  | | | | | | | |
| **TMPRSS2_1-S1/S2_2** | | | | | | | | | | | | | | | | | | | | | | | **TMPRSS2_2-S1/S2_2** | | | | | | | | | | | | | | | | | | | | |  | | | | | | | | | | | | | | | |  | | | | |  | | | | | | | | | | | | | | | |
| **Energy** | | | | | **Database** | | | | | | | **Name** | | | | | | | **Energy** | | | | | | | | | | | **Database** | | | | | | | | **Name** | | | | | |  | | | | |  | | | | | | | | | | | | | | | | | | | | | |  | | | | | | | | | |
| -12.3 | | | | | dbfda-world_531 | | | | | | | Amphotericin B | | | | | | | -12.6 | | | | | | | | | | | DrugBank_approved_980 | | | | | | | | Candicidin | | | | | |  | | | | |  | | | | | | | | | | | | | | | | | | | | | |  | | | | | | | | | |
| -12.3 | | | | | DrugBank_approved_980 | | | | | | | Candicidin | | | | | | | -12.3 | | | | | | | | | | | dbfda-world_531 | | | | | | | | Amphotericin B | | | | | |  | | | | |  | | | | | | | | | | | | | | | | | | | | | |  | | | | | | | | | |
| -12.2 | | | | | dbfda-world_496 | | | | | | | Nystatin | | | | | | | -12.1 | | | | | | | | | | | DrugBank_approved_1137 | | | | | | | | Quinupristin | | | | | |  | | | | |  | | | | | | | | | | | | | | | | | | | | | |  | | | | | | | | | |
| -11.7 | | | | | dbfda-world_494 | | | | | | | Nystatin | | | | | | | -11.8 | | | | | | | | | | | dbfda-world_256 | | | | | | | | Digoxin | | | | | |  | | | | |  | | | | | | | | | | | | | | | | | | | | | |  | | | | | | | | | |
| -11.6 | | | | | DrugBank_approved_506 | | | | | | | Nystatin | | | | | | | -11.7 | | | | | | | | | | | dbfda-world_493 | | | | | | | | Nystatin | | | | | |  | | | | |  | | | | | | | | | | | | | | | | | | | | | |  | | | | | | | | | |
| -11.5 | | | | | dbfda-world_493 | | | | | | | Nystatin | | | | | | | -11.1 | | | | | | | | | | | DrugBank_approved_506 | | | | | | | | Nystatin | | | | | |  | | | | |  | | | | | | | | | | | | | | | | | | | | | |  | | | | | | | | | |
| -11.5 | | | | | DrugBank_approved_1814 | | | | | | | Tannic acid | | | | | | | -11 | | | | | | | | | | | DrugBank_approved_2021 | | | | | | | | Velpatasvir | | | | | |  | | | | |  | | | | | | | | | | | | | | | | | | | | | |  | | | | | | | | | |
| -11.3 | | | | | dbfda-world_1534 | | | | | | | Paritaprevir | | | | | | | -11 | | | | | | | | | | | dbfda-world_494 | | | | | | | | Nystatin | | | | | |  | | | | |  | | | | | | | | | | | | | | | | | | | | | |  | | | | | | | | | |
| -11.3 | | | | | DrugBank_approved_2108 | | | | | | | Cantharidin | | | | | | | -10.9 | | | | | | | | | | | DrugBank_approved_2108 | | | | | | | | Cantharidin | | | | | |  | | | | |  | | | | | | | | | | | | | | | | | | | | | |  | | | | | | | | | |
| -11.3 | | | | | dbfda-world_1519 | | | | | | | Trypan blue | | | | | | | -10.8 | | | | | | | | | | | dbfda-world_495 | | | | | | | | Nystatin | | | | | |  | | | | |  | | | | | | | | | | | | | | | | | | | | | |  | | | | | | | | | |
| -11.2 | | | | | DrugBank_approved_2079 | | | | | | | Entrectinib | | | | | | | -10.8 | | | | | | | | | | | dbfda-world_496 | | | | | | | | Nystatin | | | | | |  | | | | |  | | | | | | | | | | | | | | | | | | | | | |  | | | | | | | | | |
| -11.2 | | | | | dbfda-world_495 | | | | | | | Nystatin | | | | | | | -10.7 | | | | | | | | | | | dbfda-world_892 | | | | | | | | Rifampicin | | | | | |  | | | | |  | | | | | | | | | | | | | | | | | | | | | |  | | | | | | | | | |
| -11.1 | | | | | DrugBank_approved_2005 | | | | | | | Venetoclax | | | | | | | -10.6 | | | | | | | | | | | dbfda-world_1136 | | | | | | | | Everolimus | | | | | |  | | | | |  | | | | | | | | | | | | | | | | | | | | | |  | | | | | | | | | |
| -11 | | | | | DrugBank_approved_3 | | | | | | | Desmopressin | | | | | | | -10.5 | | | | | | | | | | | dbfda-world_1028 | | | | | | | | Rifapentine | | | | | |  | | | | |  | | | | | | | | | | | | | | | | | | | | | |  | | | | | | | | | |
| -11 | | | | | dbfda-world_256 | | | | | | | Digoxin | | | | | | | -10.4 | | | | | | | | | | | dbfda-world_176 | | | | | | | | Dihydroergotamine | | | | | |  | | | | |  | | | | | | | | | | | | | | | | | | | | | |  | | | | | | | | | |
| -11 | | | | | dbfda-world_1028 | | | | | | | Rifapentine | | | | | | | -10.4 | | | | | | | | | | | DrugBank_approved_1639 | | | | | | | | Ledipasvir | | | | | |  | | | | |  | | | | | | | | | | | | | | | | | | | | | |  | | | | | | | | | |
| -10.9 | | | | | dbfda-world_1621 | | | | | | | Venetoclax | | | | | | | -10.3 | | | | | | | | | | | dbfda-world_690 | | | | | | | | Natamycin | | | | | |  | | | | |  | | | | | | | | | | | | | | | | | | | | | |  | | | | | | | | | |
| -10.9 | | | | | dbfda-world_690 | | | | | | | Natamycin | | | | | | | -10.3 | | | | | | | | | | | DrugBank_approved_3 | | | | | | | | Desmopressin | | | | | |  | | | | |  | | | | | | | | | | | | | | | | | | | | | |  | | | | | | | | | |
| -10.9 | | | | | TheBindingDB_660 | | | | | | | Ivacaftor | | | | | | | -10.3 | | | | | | | | | | | DrugBank_approved_2005 | | | | | | | | Venetoclax | | | | | |  | | | | |  | | | | | | | | | | | | | | | | | | | | | |  | | | | | | | | | |
| -10.8 | | | | | DrugBank_approved_1137 | | | | | | | Quinupristin | | | | | | | -10.3 | | | | | | | | | | | dbfda-world_545 | | | | | | | | Ergotamine | | | | | |  | | | | |  | | | | | | | | | | | | | | | | | | | | | |  | | | | | | | | | |
| -10.8 | | | | | TheBindingDB_770 | | | | | | | Eptifibatide | | | | | | | -10.2 | | | | | | | | | | | dbfda-world_1621 | | | | | | | | Venetoclax | | | | | |  | | | | |  | | | | | | | | | | | | | | | | | | | | | |  | | | | | | | | | |
| -10.8 | | | | | DrugBank_approved_2021 | | | | | | | Velpatasvir | | | | | | | -10.2 | | | | | | | | | | | dbfda-world_735 | | | | | | | | Sirolimus | | | | | |  | | | | |  | | | | | | | | | | | | | | | | | | | | | |  | | | | | | | | | |
| **TMPRSS2_1-S2'** | | | | | | | | | | | | | | | | | | | | | | **TMPRSS2_2-S2'** | | | | | | | | | | | | | | | | | | | | |  | | | | | | | | | | | | | | | |  | | | |  | | | | | | | | | | | | | | | | | |
| **Energy** | | | | **Database** | | | | | | | **Name** | | | | | | | **Energy** | | | | | | | | | | **Database** | | | | | | | | **Name** | | | | | | |  | | | |  | | | | | | | | | | | | | | | | | | | |  | | | | | | | | | | | | | |
| -12.1 | | | | DrugBank_approved_2108 | | | | | | | Cantharidin | | | | | | | -13.7 | | | | | | | | | | DrugBank_approved_1814 | | | | | | | | Tannic acid | | | | | | |  | | | |  | | | | | | | | | | | | | | | | | | | |  | | | | | | | | | | | | | |
| -10.9 | | | | dbfda-world_256 | | | | | | | Digoxin | | | | | | | -13.2 | | | | | | | | | | DrugBank_approved_8 | | | | | | | | Abarelix | | | | | | |  | | | |  | | | | | | | | | | | | | | | | | | | |  | | | | | | | | | | | | | |
| -10.7 | | | | ChemBridge_26 | | | | | | | 7786009 | | | | | | | -12.8 | | | | | | | | | | dbfda-world_256 | | | | | | | | Digoxin | | | | | | |  | | | |  | | | | | | | | | | | | | | | | | | | |  | | | | | | | | | | | | | |
| -10.6 | | | | DrugBank_approved_1361 | | | | | | | Indacaterol | | | | | | | -12.7 | | | | | | | | | | DrugBank_approved_2441 | | | | | | | | Ubrogepant | | | | | | |  | | | |  | | | | | | | | | | | | | | | | | | | |  | | | | | | | | | | | | | |
| -10.4 | | | | TheBindingDB_910 | | | | | | | Avagard | | | | | | | -12.6 | | | | | | | | | | dbfda-world_730 | | | | | | | | Conivaptan | | | | | | |  | | | |  | | | | | | | | | | | | | | | | | | | |  | | | | | | | | | | | | | |
| -10.4 | | | | TheBindingDB_174 | | | | | | | Raltegravir | | | | | | | -12.5 | | | | | | | | | | DrugBank_approved_2005 | | | | | | | | Venetoclax | | | | | | |  | | | |  | | | | | | | | | | | | | | | | | | | |  | | | | | | | | | | | | | |
| -10.3 | | | | TheBindingDB_87 | | | | | | | Nexavar | | | | | | | -12.3 | | | | | | | | | | dbfda-world_176 | | | | | | | | Dihydroergotamine | | | | | | |  | | | |  | | | | | | | | | | | | | | | | | | | |  | | | | | | | | | | | | | |
| -10.3 | | | | dbfda-world_494 | | | | | | | Nystatin | | | | | | | -12.2 | | | | | | | | | | DrugBank_approved_90 | | | | | | | | Adapalene | | | | | | |  | | | |  | | | | | | | | | | | | | | | | | | | |  | | | | | | | | | | | | | |
| -10.2 | | | | dbfda-world_730 | | | | | | | Conivaptan | | | | | | | -12.2 | | | | | | | | | | TheBindingDB_955 | | | | | | | | Tasigna | | | | | | |  | | | |  | | | | | | | | | | | | | | | | | | | |  | | | | | | | | | | | | | |
| -10.1 | | | | DrugBank_approved_2058 | | | | | | | Bictegravir | | | | | | | -12.1 | | | | | | | | | | TheBindingDB_1320 | | | | | | | | Tannic Acid | | | | | | |  | | | |  | | | | | | | | | | | | | | | | | | | |  | | | | | | | | | | | | | |
| -10 | | | | DrugBank_approved_1076 | | | | | | | Gliquidone | | | | | | | -12.1 | | | | | | | | | | TheBindingDB_1061 | | | | | | | | Vapsirol | | | | | | |  | | | |  | | | | | | | | | | | | | | | | | | | |  | | | | | | | | | | | | | |
| -10 | | | | DrugBank_approved_2274 | | | | | | | Netarsudil | | | | | | | -12 | | | | | | | | | | dbfda-world_75 | | | | | | | | Adapalene | | | | | | |  | | | |  | | | | | | | | | | | | | | | | | | | |  | | | | | | | | | | | | | |
| -10 | | | | TheBindingDB_1139 | | | | | | | Regorafenib | | | | | | | -12 | | | | | | | | | | DrugBank_approved_720 | | | | | | | | Conivaptan | | | | | | |  | | | |  | | | | | | | | | | | | | | | | | | | |  | | | | | | | | | | | | | |
| -10 | | | | TheBindingDB_688 | | | | | | | Irbesartan | | | | | | | -11.9 | | | | | | | | | | dbfda-world_1573 | | | | | | | | Ecamsule | | | | | | |  | | | |  | | | | | | | | | | | | | | | | | | | |  | | | | | | | | | | | | | |
| -9.9 | | | | DrugBank_approved_2030 | | | | | | | Delamanid | | | | | | | -11.9 | | | | | | | | | | DrugBank_approved_1162 | | | | | | | | Antrafenine | | | | | | |  | | | |  | | | | | | | | | | | | | | | | | | | |  | | | | | | | | | | | | | |
| -9.9 | | | | DrugBank_approved_1694 | | | | | | | Daclatasvir | | | | | | | -11.9 | | | | | | | | | | TheBindingDB_408 | | | | | | | | Vapsirol | | | | | | |  | | | |  | | | | | | | | | | | | | | | | | | | |  | | | | | | | | | | | | | |
| -9.9 | | | | DrugBank_approved_377 | | | | | | | Acetyldigitoxin | | | | | | | -11.9 | | | | | | | | | | TheBindingDB_402 | | | | | | | | Synarel | | | | | | |  | | | |  | | | | | | | | | | | | | | | | | | | |  | | | | | | | | | | | | | |
| -9.9 | | | | ChemBridge_10 | | | | | | | 7793889 | | | | | | | -11.9 | | | | | | | | | | TheBindingDB_772 | | | | | | | | Tolvaptan | | | | | | |  | | | |  | | | | | | | | | | | | | | | | | | | |  | | | | | | | | | | | | | |
| -9.8 | | | | dbfda-world_1085 | | | | | | | Paliperidone | | | | | | | -11.8 | | | | | | | | | | dbfda-world_1455 | | | | | | | | Ledipasvir | | | | | | |  | | | |  | | | | | | | | | | | | | | | | | | | |  | | | | | | | | | | | | | |
| -9.9 | | | | **-** | | | | | | | Camostat | | | | | | | -11.8 | | | | | | | | | | dbfda-world_1396 | | | | | | | | Lomitapide | | | | | | |  | | | | |  | | | | | | | | | | | | | | | | | | | | |  | | | | | | | | | | | |
| **catL_1-S1/S2’_1^+^** | | | | | | | | | | | | | | | | | | | | | | **catL_2-S1/S2’_1^+^** | | | | | | | | | | | | | | | | | | | | | **catL_3-S1/S2’_1^+^** | | | | | | | | | | | | | | | | | | | | | | | | | | | | | | | | | | | | | |
| **Energy** | | | | **Database** | | | | | | | **Name** | | | | | | | **Energy** | | | | | | | | | | **Database** | | | | | | | | **Name** | | | | | | | **Energy** | | | | | | | | | **Database** | | | | | | | | | | | | | | | | | | | | | | | | | **Name** | | | |
| -11.1 | | | | DrugBank_approved_980 | | | | | | | Candicidin | | | | | | | -11.9 | | | | | | | | | | dbfda-world_1572 | | | | | | | | Ecamsule | | | | | | | -11.4 | | | | | | | | | dbfda-world_531 | | | | | | | | | | | | | | | | | | | | | | | | | Amphotericin B | | | |
| -11 | | | | dbfda-world_494 | | | | | | | Nystatin | | | | | | | -11.8 | | | | | | | | | | dbfda-world_1573 | | | | | | | | Ecamsule | | | | | | | -11.3 | | | | | | | | | dbfda-world_496 | | | | | | | | | | | | | | | | | | | | | | | | | Nystatin | | | |
| -10.6 | | | | dbfda-world_256 | | | | | | | Digoxin | | | | | | | -11.4 | | | | | | | | | | dbfda-world_632 | | | | | | | | Irinotecan | | | | | | | -11.2 | | | | | | | | | dbfda-world_256 | | | | | | | | | | | | | | | | | | | | | | | | | Digoxin | | | |
| -10.4 | | | | DrugBank_approved_1137 | | | | | | | Quinupristin | | | | | | | -11.4 | | | | | | | | | | DrugBank_approved_2108 | | | | | | | | Cantharidin | | | | | | | -10.7 | | | | | | | | | DrugBank_approved_1137 | | | | | | | | | | | | | | | | | | | | | | | | | Quinupristin | | | |
| -10.2 | | | | DrugBank_approved_506 | | | | | | | Nystatin | | | | | | | -11.4 | | | | | | | | | | dbfda-world_256 | | | | | | | | Digoxin | | | | | | | -10.7 | | | | | | | | | dbfda-world_495 | | | | | | | | | | | | | | | | | | | | | | | | | Nystatin | | | |
| -10.2 | | | | dbfda-world_495 | | | | | | | Nystatin | | | | | | | -11.3 | | | | | | | | | | dbfda-world_176 | | | | | | | | Dihydroergotamine | | | | | | | -10.6 | | | | | | | | | DrugBank_approved_506 | | | | | | | | | | | | | | | | | | | | | | | | | Nystatin | | | |
| -10.1 | | | | DrugBank_approved_3 | | | | | | | Desmopressin | | | | | | | -11.2 | | | | | | | | | | dbfda-world_730 | | | | | | | | Conivaptan | | | | | | | -10.4 | | | | | | | | | DrugBank_approved_540 | | | | | | | | | | | | | | | | | | | | | | | | | Amphotericin B | | | |
| -10.1 | | | | dbfda-world_496 | | | | | | | Nystatin | | | | | | | -11.2 | | | | | | | | | | dbfda-world_1428 | | | | | | | | Ponatinib | | | | | | | -10.3 | | | | | | | | | DrugBank_approved_980 | | | | | | | | | | | | | | | | | | | | | | | | | Candicidin | | | |
| -10 | | | | dbfda-world_531 | | | | | | | Amphotericin B | | | | | | | -11.2 | | | | | | | | | | dbfda-world_1519 | | | | | | | | Trypan blue | | | | | | | -10.3 | | | | | | | | | dbfda-world_494 | | | | | | | | | | | | | | | | | | | | | | | | | Nystatin | | | |
| -10 | | | | dbfda-world_545 | | | | | | | Ergotamine | | | | | | | -11.2 | | | | | | | | | | TheBindingDB_511 | | | | | | | | Accolate | | | | | | | -10.2 | | | | | | | | | DrugBank_approved_1401 | | | | | | | | | | | | | | | | | | | | | | | | | Icatibant | | | |
| -9.8 | | | | dbfda-world_176 | | | | | | | Dihydroergotamine | | | | | | | -11.1 | | | | | | | | | | DrugBank_approved_2079 | | | | | | | | Entrectinib | | | | | | | -10.1 | | | | | | | | | DrugBank_approved_3 | | | | | | | | | | | | | | | | | | | | | | | | | Desmopressin | | | |
| -9.8 | | | | dbfda-world_1634 | | | | | | | Deflazacort | | | | | | | -11.1 | | | | | | | | | | dbfda-world_1263 | | | | | | | | Eltrombopag | | | | | | | -10.1 | | | | | | | | | DrugBank_approved_1639 | | | | | | | | | | | | | | | | | | | | | | | | | Ledipasvir | | | |
| -9.8 | | | | DrugBank_approved_2005 | | | | | | | Venetoclax | | | | | | | -11 | | | | | | | | | | dbfda-world_1265 | | | | | | | | Tolvaptan | | | | | | | -10 | | | | | | | | | DrugBank_approved_1339 | | | | | | | | | | | | | | | | | | | | | | | | | Nilotinib | | | |
| -9.8 | | | | dbfda-world_1028 | | | | | | | Rifapentine | | | | | | | -10.9 | | | | | | | | | | DrugBank_approved_555 | | | | | | | | Ergotamine | | | | | | | -9.9 | | | | | | | | | DrugBank_approved_555 | | | | | | | | | | | | | | | | | | | | | | | | | Ergotamine | | | |
| -9.7 | | | | dbfda-world_1534 | | | | | | | Paritaprevir | | | | | | | -10.9 | | | | | | | | | | dbfda-world_1425 | | | | | | | | Regorafenib | | | | | | | -9.9 | | | | | | | | | TheBindingDB_955 | | | | | | | | | | | | | | | | | | | | | | | | | Tasigna | | | |
| -9.7 | | | | DrugBank_approved_811 | | | | | | | Dactinomycin | | | | | | | -10.8 | | | | | | | | | | dbfda-world_1075 | | | | | | | | Retapamulin | | | | | | | -9.9 | | | | | | | | | TheBindingDB_1061 | | | | | | | | | | | | | | | | | | | | | | | | | Vapsirol | | | |
| -9.7 | | | | DrugBank_approved_2108 | | | | | | | Cantharidin | | | | | | | -10.8 | | | | | | | | | | DrugBank_approved_1670 | | | | | | | | Edoxaban | | | | | | | -9.9 | | | | | | | | | DrugBank_approved_1046 | | | | | | | | | | | | | | | | | | | | | | | | | Rifaximin | | | |
| -9.6 | | | | TheBindingDB_435 | | | | | | | Fluorometholone | | | | | | | -10.8 | | | | | | | | | | DrugBank_approved_720 | | | | | | | | Conivaptan | | | | | | | -9.8 | | | | | | | | | TheBindingDB_408 | | | | | | | | | | | | | | | | | | | | | | | | | Vapsirol | | | |
| -9.6 | | | | DrugBank_approved_555 | | | | | | | Ergotamine | | | | | | | -10.7 | | | | | | | | | | DrugBank_approved_125 | | | | | | | | Ziprasidone | | | | | | | -9.8 | | | | | | | | | dbfda-world_493 | | | | | | | | | | | | | | | | | | | | | | | | | Nystatin | | | |
|  | | | |  | | | | | | |  | | | | | | | -10.7 | | | | | | | | | | DrugBank_approved_2020 | | | | | | | | Lifitegrast | | | | | | | -9.7 | | | | | | | | | DrugBank_approved_2079 | | | | | | | | | | | | | | | | | | | | | | | | | Entrectinib | | | |
|  | | | |  | | | | | | |  | | | | | | | -10.7 | | | | | | | | | | DrugBank_approved_2005 | | | | | | | | Venetoclax | | | | | | | -9.7 | | | | | | | | | DrugBank_approved_1814 | | | | | | | | | | | | | | | | | | | | | | | | | Tannic acid | | | |
| **catL_1-S1/S2’_2^+^** | | | | | | | | | | | | | | | | | | | | | | **catL_2-S1/S2’_2^+^** | | | | | | | | | | | | | | | | | | | | | **catL_3-S1/S2’_2^+^** | | | | | | | | | | | | | | | | | | | | | | | | | | | | | | | | | | | | | |
| **Energy** | | | | | | **Database** | | | | | | | **Name** | | | | | | | | | | | **Energy** | | | | | | | **Database** | | | | | | | | **Name** | | | | **Energy** | | | | | | | | | | | **Database** | | | | | | | | | | | | | | | | | | | | | | | | | **Name** | |
| -11.1 | | | | | | dbfda-world_531 | | | | | | | Amphotericin B | | | | | | | | | | | -11.4 | | | | | | | dbfda-world_1263 | | | | | | | | Eltrombopag | | | | -12 | | | | | | | | | | | dbfda-world_494 | | | | | | | | | | | | | | | | | | | | | | | | | Nystatin | |
| -11.1 | | | | | | dbfda-world_256 | | | | | | | Digoxin | | | | | | | | | | | -11.3 | | | | | | | dbfda-world_256 | | | | | | | | Digoxin | | | | -11.2 | | | | | | | | | | | dbfda-world_531 | | | | | | | | | | | | | | | | | | | | | | | | | Amphotericin B | |
| -10.6 | | | | | | dbfda-world_493 | | | | | | | Nystatin | | | | | | | | | | | -11.1 | | | | | | | dbfda-world_730 | | | | | | | | Conivaptan | | | | -11 | | | | | | | | | | | dbfda-world_495 | | | | | | | | | | | | | | | | | | | | | | | | | Nystatin | |
| -10.3 | | | | | | dbfda-world_1519 | | | | | | | Trypan blue | | | | | | | | | | | -10.9 | | | | | | | TheBindingDB_780 | | | | | | | | Juxtapid | | | | -10.9 | | | | | | | | | | | DrugBank_approved_1137 | | | | | | | | | | | | | | | | | | | | | | | | | Quinupristin | |
| -10.3 | | | | | | DrugBank_approved_980 | | | | | | | Candicidin | | | | | | | | | | | -10.7 | | | | | | | dbfda-world_1629 | | | | | | | | Velpatasvir | | | | -10.9 | | | | | | | | | | | DrugBank_approved_980 | | | | | | | | | | | | | | | | | | | | | | | | | Candicidin | |
| -10.2 | | | | | | DrugBank_approved_1137 | | | | | | | Quinupristin | | | | | | | | | | | -10.7 | | | | | | | TheBindingDB_955 | | | | | | | | Tasigna | | | | -10.7 | | | | | | | | | | | dbfda-world_1048 | | | | | | | | | | | | | | | | | | | | | | | | | Rifaximin | |
| -10.2 | | | | | | dbfda-world_176 | | | | | | | Dihydroergotamine | | | | | | | | | | | -10.7 | | | | | | | dbfda-world_1534 | | | | | | | | Paritaprevir | | | | -10.7 | | | | | | | | | | | DrugBank_approved_725 | | | | | | | | | | | | | | | | | | | | | | | | | Sirolimus | |
| -10.1 | | | | | | dbfda-world_1534 | | | | | | | Paritaprevir | | | | | | | | | | | -10.6 | | | | | | | DrugBank_approved_1409 | | | | | | | | Eltrombopag | | | | -10.5 | | | | | | | | | | | DrugBank_approved_2026 | | | | | | | | | | | | | | | | | | | | | | | | | Temoporfin | |
| -10.1 | | | | | | dbfda-world_495 | | | | | | | Nystatin | | | | | | | | | | | -10.6 | | | | | | | dbfda-world_496 | | | | | | | | Nystatin | | | | -10.5 | | | | | | | | | | | dbfda-world_496 | | | | | | | | | | | | | | | | | | | | | | | | | Nystatin | |
| -10.1 | | | | | | dbfda-world_494 | | | | | | | Nystatin | | | | | | | | | | | -10.5 | | | | | | | DrugBank_approved_2079 | | | | | | | | Entrectinib | | | | -10.5 | | | | | | | | | | | dbfda-world_256 | | | | | | | | | | | | | | | | | | | | | | | | | Digoxin | |
| -10 | | | | | | DrugBank_approved_1814 | | | | | | | Tannic acid | | | | | | | | | | | -10.5 | | | | | | | TheBindingDB_989 | | | | | | | | Promacta | | | | -10.4 | | | | | | | | | | | DrugBank_approved_555 | | | | | | | | | | | | | | | | | | | | | | | | | Ergotamine | |
| -9.9 | | | | | | DrugBank_approved_3 | | | | | | | Desmopressin | | | | | | | | | | | -10.5 | | | | | | | dbfda-world_494 | | | | | | | | Nystatin | | | | -10.4 | | | | | | | | | | | dbfda-world_1534 | | | | | | | | | | | | | | | | | | | | | | | | | Paritaprevir | |
| -9.8 | | | | | | DrugBank_approved_2265 | | | | | | | Pibrentasvir | | | | | | | | | | | -10.4 | | | | | | | dbfda-world_495 | | | | | | | | Nystatin | | | | -10.4 | | | | | | | | | | | dbfda-world_493 | | | | | | | | | | | | | | | | | | | | | | | | | Nystatin | |
| -9.8 | | | | | | dbfda-world_1621 | | | | | | | Venetoclax | | | | | | | | | | | -10.4 | | | | | | | DrugBank_approved_1639 | | | | | | | | Ledipasvir | | | | -10.3 | | | | | | | | | | | dbfda-world_1654 | | | | | | | | | | | | | | | | | | | | | | | | | Glecaprevir | |
| -9.8 | | | | | | DrugBank_approved_377 | | | | | | | Acetyldigitoxin | | | | | | | | | | | -10.4 | | | | | | | DrugBank_approved_2005 | | | | | | | | Venetoclax | | | | -10.3 | | | | | | | | | | | dbfda-world_1299 | | | | | | | | | | | | | | | | | | | | | | | | | Midostaurin | |
| -9.8 | | | | | | dbfda-world_496 | | | | | | | Nystatin | | | | | | | | | | | -10.3 | | | | | | | DrugBank_approved_1814 | | | | | | | | Tannic acid | | | | -10.2 | | | | | | | | | | | DrugBank_approved_3 | | | | | | | | | | | | | | | | | | | | | | | | | Desmopressin | |
| -9.8 | | | | | | DrugBank_approved_1543 | | | | | | | Triptorelin | | | | | | | | | | | -10.3 | | | | | | | DrugBank_approved_1401 | | | | | | | | Icatibant | | | | -10.2 | | | | | | | | | | | DrugBank_approved_1639 | | | | | | | | | | | | | | | | | | | | | | | | | Ledipasvir | |
| -9.8 | | | | | | DrugBank_approved_2108 | | | | | | | Cantharidin | | | | | | | | | | | -10.3 | | | | | | | dbfda-world_1529 | | | | | | | | Lumacaftor | | | | -10.2 | | | | | | | | | | | dbfda-world_1519 | | | | | | | | | | | | | | | | | | | | | | | | | Trypan blue | |
| -9.8 | | | | | | dbfda-world_1028 | | | | | | | Rifapentine | | | | | | | | | | | -10.3 | | | | | | | DrugBank_approved_479 | | | | | | | | Imatinib | | | | -10.1 | | | | | | | | | | | dbfda-world_176 | | | | | | | | | | | | | | | | | | | | | | | | | Dihydroergotamine | |
| -9.7 | | | | | | dbfda-world_1572 | | | | | | | Ecamsule | | | | | | | | | | | -10.3 | | | | | | | DrugBank_approved_2108 | | | | | | | | Cantharidin | | | | -10.1 | | | | | | | | | | | dbfda-world_736 | | | | | | | | | | | | | | | | | | | | | | | | | Sirolimus | |
|  | | | | | |  | | | | | | |  | | | | | | | | | | |  | | | | | | |  | | | | | | | |  | | | | -10 | | | | | | | | | | | dbfda-world_1027 | | | | | | | | | | | | | | | | | | | | | | | | | Bromocriptine | |

*Unit of energy is kcal/mol. AutoDock Vina (version 1.1.2, Linux) was used for the docking.

+ S1/S2’ corresponds to the cleavage site targeted by catL (T696-M697) on S1/S2 boundary.

**Table S3.** The average binding free energy of protein-ligand complexes*

| **Complex** | $\boldsymbol{\Delta G}$ |  | **Complex** | $\boldsymbol{\Delta G}$ |  | **Complex** | $\boldsymbol{\Delta G}$ |
| --- | --- | --- | --- | --- | --- | --- | --- |
| trypsin-S1/S2-Digoxin | -189.21922 |  | TMPRSS2-S2'-Ubrogepant | -44.6749 |  | catL-Digoxin | 468.1419 |
| trypsin-S1/S2-FAD | -186.88962 |  | S1/S2-Ciclesonide | -40.2 |  | catL-S1/S2'-Irinotecan | 469.175 |
| S1/S2-Avatrombopag | -154.4893 |  | trypsin-Digoxin | -22.3252 |  | catL-S1/S2'-Vapsirol | 477.135 |
| trypsin-S2'-Ursodeaoxycholic acid | -154.29806 |  | catL-Nystatin | -17.5213 |  | S2'-D936Y-Saquinavir | 490.7611 |
| TMPRSS2-S1/S2-Digoxin | -147.1056 |  | TMPRSS2-Nystatin | -8.97008 |  | trypsin-Dutasteride | 545.7077 |
| catL-S1/S2'-Rifapentine | -138.49152 |  | S1/S2-Nystatin | -4.07952 |  | S1/S2-Rifaximin | 574.1273 |
| trypsin-S1/S2-Adapalene | -121.2825 |  | trypsin-S2'-Dihydroergotamine | 0.9803 |  | trypsin-S1/S2-Bictegravir | 579.2934 |
| TMPRSS2-S1/S2-Telmisartan | -121.09444 |  | S1/S2-Capmatinib | 29.65224 |  | trypsin-Digoxin | 594.7297 |
| catL-Digoxin | -120.32792 |  | S1/S2'-Ubrogepant | 31.30914 |  | S1/S2-Ubrogepant | 608.25448 |
| S2'-Antrafenine | -118.58336 |  | catL-S1/S2'-Nystatin | 40.80676 |  | trypsin-S2'-Digoxin | 608.7868 |
| TMPRSS2-S1/S2-Digoxin | -116.02408 |  | catL-Digoxin | 45.23074 |  | trypsin-S1/S2-Dihydroergotamine | 613.8832 |
| catL-S1/S2'-Rifapentine | -114.56038 |  | trypsin-Valrubicin | 52.70146 |  | S1/S2'-Nystatin | 621.6618 |
| trypsin-S2'-Lasmiditan | -109.25438 |  | TMPRSS2-Sirolimus | 67.8708 |  | t1-S2'-D936Y-Nebivolol | 640.6687 |
| trypsin-S2'-Glimepiride | -107.87552 |  | TMPRSS2-Digoxin | 74.04584 |  | TMPRSS2-S2'-Dihydroergotamine | 641.7708 |
| TMPRSS2-S2'-Bictegravir | -104.40558 |  | TMPRSS2-Drospirenone | 83.41938 |  | trypsin-S1/S2-Dihydroergotamine | 644.3816 |
| catL-Drospirenone | -101.37212 |  | catL-Rifaximin | 91.25842 |  | TMPRSS2-S2'-Digoxin | 647.4213 |
| TMPRSS2-Dexamethasone M | -99.6151 |  | S2'-D936Y-Glyxambi | 95.0251 |  | TMPRSS2-Ubrogepant | 657.21852 |
| trypsin-S1/S2-Ritonavir | -99.27076 |  | trypsin-Digoxin | 109.0991 |  | trypsin-S2'-Nystatin | 710.1351 |
| S2'-D936Y-Irbesartan | -95.90374 |  | trypsin-S1/S2-Droperidol | 110.7961 |  | catL-S1/S2'-Edoxaban | 714.1895 |
| TMPRSS2-Digoxin | -95.62794 |  | S1/S2-Vapsirol | 111.637 |  | catL-S1/S2'-Digoxin | 719.4035 |
| S2'-D936Y-Nebivolol | -94.06474 |  | catL-S1/S2'-Pibrentasvir | 111.949 |  | trypsin-S2'-Aprepitant | 725.1868 |
| trypsin-S1/S2-Drospirenone | -93.28244 |  | S2'-D936Y-Deoxycholic acid | 113.3931 |  | catL-S1/S2'-Drospirenone | 736.8676 |
| catL-S1/S2'-Deflazacort | -91.99702 |  | S2'-D936Y-Viracept | 133.7918 |  | catL-Dihydroergotamine | 817.161 |
| TMPRSS2-S2'-Gliquidone | -90.84554 |  | trypsin-S1/S2-Dihydroergotamine | 143.2903 |  | trypsin-S1/S2-Dihydroergotamine | 845.0419 |
| TMPRSS2-S2'-Antrafenine | -90.32412 |  | trypsin-S1/S2-Antrafenine | 151.7891 |  | trypsin-S1/S2-Gleevec | 848.3642 |
| TMPRSS2-S1/S2-Nystatin | -87.9875 |  | TMPRSS2-S1/S2-Elbasvir | 152.1473 |  | S1/S2-Digoxin | 870.507 |
| S1/S2'-Digoxin | -86.89394 |  | trypsin-S1/S2-Capmatinib | 170.6533 |  | S2'-Digoxin | 891.34192 |
| catL-S1/S2'-Rifaximin | -84.2909 |  | catL-S1/S2'-Dihydroergotamine | 170.7121 |  | trypsin-S2'-Antrafenine | 915.6934 |
| TMPRSS2-Ubrogepant | -83.68272 |  | TMPRSS2-S1/S2-Dihydroergotamine | 176.0496 |  | t2-S2'-D936Y-Atovaquone | 991.0772 |
| catL-S1/S2'-Digoxin | -83.00332 |  | S1/S2-Dihydroergotamine | 178.0787 |  | trypsin-Dihydroergotamine | 994.1542 |
| trypsin-S1/S2-Digoxin | -81.86608 |  | catL-S1/S2'-Fluorop | 187.3583 |  | TMPRSS2-S2'-Daclatasvir | 1050.959 |
| S1/S2-Cefpiramide | -81.200879 |  | catL-S1/S2'-Dihydroergotamine | 200.0495 |  | catL-Dihydroergotamine | 1071.849 |
| catL-Rifapentine | -81.06458 |  | TMPRSS2-S2'-Adapalene | 215.1768 |  | trypsin-Saquinavir | 1113.378 |
| trypsin-Saquinavir | -80.90818 |  | t2-S2'-D936Y-Paliperidone | 215.225959 |  | TMPRSS2-S1/S2-Drospirenone | 1120.802 |
| S1/S2'-Rifapentine | -80.62622 |  | t1-S2'-D936Y-Rolapitant | 228.956 |  | TMPRSS2-S1/S2-Sirolimus | 1123.432 |
| trypsin-S1/S2-Capmatinib | -75.69074 |  | trypsin-S2'-Raltegravir | 229.0492 |  | catL-S1/S2'-Dihydroergotamine | 1150.167 |
| TMPRSS2-S1/S2-Nystatin | -74.57538 |  | trypsin-S2'-Axinitib | 229.4983 |  | S1/S2-Digoxin | 1252.084 |
| S1/S2-Digoxin | -74.53658 |  | trypsin-S2'-Vapsirol | 230.2532 |  | S2'-Drospirenone | 1339.57034 |
| TMPRSS2-Saquinavir | -73.14612 |  | trypsin-S1/S2-Nystatin | 233.5296 |  | S1/S2'-Drospirenone | 1371.384 |
| S1/S2'-Rifaximin | -71.57796 |  | TMPRSS2-Dihydroergotamine | 237.2782 |  | trypsin-S1/S2-Vemurafenib | 1392.289 |
| trypsin-Drospirenone | -70.2387 |  | trypsin-S2'-Vapsirol | 247.7259 |  | TMPRSS2-S2'-Digoxin | 1411.963 |
| S1/S2'-Dihydroergotamine | -69.52424 |  | TMPRSS2-S1/S2-Simeprevir | 253.0695 |  | catL-S1/S2'-Digoxin | 1543.352 |
| S1/S2'-Capmatinib | -68.23316 |  | S1/S2-Vapsirol | 269.363 |  | catL-S1/S2'-Nystatin | 1575.642 |
| t2-S2'-D936Y-Risperidone | -67.53934 |  | TMPRSS2-Vapsirol | 272.8234 |  | trypsin-S2'-Elbasvir | 1634.338 |
| S1/S2'-Digoxin | -67.3627 |  | trypsin-S2'-Digoxin | 286.9939 |  | TMPRSS2-S2'-Delamanid | 1704.878 |
| trypsin-Rifapentine | -67.03428 |  | S2'-D936Y-Atovaquone | 298.3715 |  | trypsin-S2'-Viibryd | 1725.864 |
| trypsin-S1/S2-Ubrogepant | -64.71386 |  | catL-Dihydroergotamine | 311.9322 |  | TMPRSS2-S2'-Irbesartan | 1757.756 |
| trypsin-S1/S2-Digoxin | -63.72308 |  | catL-S1/S2'-Promacta | 313.5024 |  | trypsin-S1/S2-Rifapentine | 1810.328 |
| TMPRSS2-S2'-Vapsirol | -63.36442 |  | trypsin-S1/S2-Digoxin | 385.2007 |  | TMPRSS2-S1/S2-Dihydroergotamine | 2025.632 |
| TMPRSS2-S1/S2-Nystatin | -63.1792 |  | TMPRSS2-Dexamethasone M | 388.51754 |  | catL-S1/S2'-Digoxin | 2417.164 |
| t1-S2'-D936Y-Olaparib | -62.88132 |  | TMPRSS2-S1/S2-Rifapentine | 390.1107 |  | S1/S2'-Digoxin | 2422.388 |
| trypsin-S2'-Prelay | -61.45806 |  | trypsin-S1/S2-Rifapentine | 405.8055 |  | trypsin-S1/S2-Digoxin | 2458.327 |
| catL-S1/S2'-Nystatin | -61.40966 |  | trypsin-S1/S2-Ilevro | 408.7774 |  | TMPRSS2-S2'-Raltegravir | 2485.941 |
| TMPRSS2-S1/S2-Dihydroergotamine | -61.17566 |  | trypsin-S1/S2-Diosmin | 420.0925 |  | TMPRSS2-S2'-Indacaterol | 2553.111 |
| trypsin-S1/S2-Nystatin | -54.27652 |  | trypsin-S1/S2-Ibrutinib | 423.8819 |  | TMPRSS2-S1/S2-Rifapentine | 2589.211 |
| trypsin-S1/S2-Dihydroergotamine | -51.3636 |  | TMPRSS2-S1/S2-Digoxin | 426.7328 |  | trypsin-Nystatin | 3320.98 |
| catL-S1/S2'-Dihydroergotamine | -49.38546 |  | TMPRSS2-S1/S2-Digoxin | 463.0654 |  | TMPRSS2-Digoxin | 4537.587 |
| S1/S2-Dihydroergotamine | -46.31082 |  | catL-S1/S2'-Tolvaptan | 463.8297 |  |  |  |

*Unit of $\Delta G$is kJ/mol.
